# NSC95397 is a Novel HIV-1 Latency Reversing Agent

**DOI:** 10.1101/2023.05.24.542213

**Authors:** Randilea Nichols Doyle, Vivian Yang, Yetunde I. Kayode, Robert Damoiseaux, Harry E. Taylor, Oliver I. Fregoso

## Abstract

The latent viral reservoir represents one of the major barriers of curing HIV-1. Focus on the “kick and kill” approach, in which virus expression is reactivated then cells producing virus are selectively depleted, has led to the discovery of many latency reversing agents (LRAs) that have furthered our understanding of the mechanisms driving HIV-1 latency and latency reversal. Thus far, individual compounds have yet to be robust enough to work as a therapy, highlighting the importance of identifying new compounds that target novel pathways and synergize with known LRAs. In this study, we identified a promising LRA, NSC95397, from a screen of ∼4250 compounds. We validated that NSC95397 reactivates latent viral transcription and protein expression from cells with unique integration events and across different latency models. Co-treating cells with NSC95397 and known LRAs demonstrated that NSC95397 synergizes with different drugs under both standard normoxic and physiological hypoxic conditions. NSC95397 does not globally increase open chromatin, and bulk RNA sequencing revealed NSC95397 does not greatly increase cellular transcription. Instead, NSC95397 downregulates pathways key to metabolism, cell growth, and DNA repair – highlighting the potential of these pathways in regulating HIV-1 latency. Overall, we identified NSC95397 as a novel LRA that does not largely alter global transcription, that shows potential for synergy with known LRAs, and that may act through novel pathways not previously recognized for their ability to modulate HIV-1 latency.

**Importance:** One of the largest barriers to curing HIV-1 is the latent viral reservoir – this is when the virus incorporates itself into long-lived cells in the body, ready to reactivate and re-seed infection. Destroying dormant HIV-1 is one potential pathway to a cure, yet no therapeutics have been discovered to work well in patients. In our study, we identified a compound, NSC95397, that can awaken dormant HIV-1 on its own through novel mechanisms not previously linked to HIV-1 latency. Moreover, NSC95397 improves the abilities of previously identified compounds to reactivate latent HIV-1. Thus, our study has identified a compound that can help towards the better understanding of an HIV-1 cure.

## Introduction

As antiretroviral therapy (ART) has successfully prolonged the life expectancy of people living with HIV (PLWH), these individuals are now at higher risk from comorbidities such as cancers and HIV-associated neurocognitive disorders. Latency reversal remains one of the primary hopes of a cure for PLWH. However, our lack of understanding about the molecular mechanisms leading to latency and reactivation has impeded our capacity to efficiently reactivate latent HIV-1. While some components of latency are inherent to the virus, such as the expression of Tat and cis-regulatory elements encoded within HIV-1, it seems that the primary determinants of latency and reactivation are host cellular factors that control gene expression (1, 2). Once integrated, expression of the HIV-1 provirus is subject to much of the same cellular machinery that regulates host genes. The HIV-1 LTR has multiple binding sites for the transcription factor NF-κB, as well as binding sites for both constitutive (Sp1) and inducible (NF-AT and AP-1) transcription factors (2, 3). Since transcription is dependent on the accessibility of these transcription factor binding sites, histone positioning and epigenetic modifications have large effects on HIV-1 gene expression (1, 2). Consistent with this data, the most promising latency reversing agents (LRAs) attempt to activate host transcription (such as PKC, JNK/MAPK, and NF-κB agonists) and chromatin decondensation (such as histone deacetylase inhibitors, histone methyltransferase inhibitors, and DNA methyltransferase inhibitors). PKC agonists, such as PMA and prostratin, can potently activate latent virus in cell lines but have not been taken to clinical trials for various reasons, including oncogenic potential (4). Similarly, other compounds targeting these pathways, such as Bryostatin-1 and SAHA (vorinostat), performed well *in vitro* and *ex vivo* but had limited success in clinical trials (4, 5). The likely reason for the lack of viable LRAs is the heterogeneity of the viral reservoir *in vivo* (5), as well as difficulty modeling physiological conditions *in vitro* (6). To overcome this, new LRAs are needed, especially ones acting through different mechanisms and that reactivate HIV-1 expression in different cellular conditions. At the same time, mechanistically distinct LRAs have the potential to synergize and increase the desired effect of reactivating latent virus with lower concentrations of drug and thus minimizing side effects. Future treatments will likely need to be combinatorial to overcome toxic side effects as well as stimulate as many latent cells as possible.

To address the need for LRAs stimulating different pathways and mechanisms than current ones, we performed a medium-throughput screen in J-Lat 10.6 cells, a clonally selected Jurkat cell line with an integrated ΔEnv HIV-1 that contains GFP in the place of Nef (7-9). At steady state, these cells express little to no HIV-1 proteins or GFP, but upon activation with a LRA, such as PMA, HIV-1 proteins and GFP are expressed. We started with approximately 4,250 unique small molecules that we narrowed down through multiple steps of validation, including flow cytometry, qPCR, and western blot for both HIV-1 and GFP, to a single compound, NSC95397.

Previously studied for its potential to treat cancer, NSC95397 is a Cdc25 phosphatase inhibitor that can also inhibit many members of the dual-specificity phosphatase (DUSP) family, specifically DUSP1 (MKP1), DUSP6 (MKP3), and DSUP14 (MKP6) (10-13), which are key to cell proliferation and reaction to infection. NSC95397 has been primarily studied in cancer cells, with a focus on its ability to inhibit cell growth and induce cell cycle arrest (11). Only a few studies have been conducted in immune cells, where NSC95397 is suggested to be anti-inflammatory through inhibition of serine-threonine kinases such CD45 and SHP-2 (14). Additionally, most transcription factor families associated with HIV-1 transcription are inhibited in NSC95397 treated cells (15), making the mechanism by which NSC95397 activates HIV-1 transcription unclear.

To understand how NSC95397 reactivates latent HIV-1, we first tested whether NSC95397 synergizes with known LRAs. We performed combinatorial studies with various LRAs and found that NSC95397 has additive and synergistic effects on reactivation when combined with the tested LRAs. The synergistic effects were further maintained in the ACH-2 T-cell line (a polyclonal cell line infected with Tat/tar functionally deficient HIV-1, resulting in different integrations sites as well as low levels of active HIV-1 replication) (16-18), and the U1 monocytic line (a polyclonal cell line infected with Tat-deficient HIV-1, resulting in different integrations sites but minimal background replication) (18, 19). Analysis of histone marks associated with active transcription indicated that NSC95397 does not alter global histone modifications to reactivate latent HIV-1. Transcriptome analysis by RNA sequencing showed that NSC95397 also does not stimulate transcriptional upregulation or Tat-mediated LTR activation. Instead, NSC95397 leads to the down-regulation of diverse pathways, including metabolism, cell cycle, hypoxia/redox, and DNA repair. We confirmed that cells treated with NSC95397 have increased accumulation of signaling molecules in the DNA damage repair pathways. To test the efficacy of NSC95397 under conditions found in hypoxic tissues that harbor the HIV-1 reservoir *in vivo*, we also assessed activity under physiological hypoxia (20). NSC95397 reactivates latent HIV-1 and maintains its ability to increase reactivation in combination with other LRAs even under hypoxia. Thus, while we were unable to identify a definitive mechanism of action for NSC95397, our data using multiple models of HIV-1 latency indicate that NSC95397 acts on different and unknown pathway(s) compared to current LRAs and could be a useful tool in understanding the molecular mechanisms of HIV-1 latency and as combinatorial therapy in our quest to cure latent HIV-1.

## Results

### NSC95397 identified as an HIV-1 latency reversal agent (LRA) from a medium-throughput screen

To identify novel compounds and molecular pathways of HIV-1 latency reversal, we conducted a medium-throughput screen of approximately 4,250 compounds (Figure 1A) in J-Lat 10.6 cells, a clonally selected Jurkat cell line with a latently integrated ΔEnv HIV-1 that contains GFP in the place of Nef (7, 8). In the initial screen, we incubated the J-Lat 10.6 cells with individual compounds in 348-well glass-bottom plates for 36 hours, then imaged for GFP expression using an automated ImageExpress microscope (Figure S1A-B). Using z-score cutoffs of ≥ 2 for GFP cell count and % of GFP positive cells and ≥ 1 for GFP average intensity, we identified 83 compounds for additional testing (Figure 1B-C). Though PMA was a positive control used in this study, we also found it as a hit within the library (light grey square in Figure 1B-C, S1A-B). Additionally, we identified IBET151 (purple diamond in Figure 1B-C, S1A-B), a previously described LRA (21), suggesting our screen is robust enough to identify new potential LRAs. To validate the 83 hits, GFP expression was measured in J-Lat 10.6 cells treated in triplicate and incubated for 24 and 48 hours (Figure 1D). Thirty-five of the 83 compounds validated with repeated GFP expression. Neuronal signaling, DNA damage response (DDR) pathways, and metabolism were the top three pathways targeted by the 35 validated compounds (Figure 1E). Next, we performed EC_50_ titration curves on a subset of the validated compounds in the J-Lat 10.6 and 5A8 cells, another clonal J-Lat line with a unique integration event (9), at 24 and 48 hours (Figure S1C-E). NSC95397 further validated at multiple concentrations in both 10.6 and 5A8 J-Lat cells with little toxicity (Figure 1F; blue triangle in Figure 1B-C, S1). Unlike NSC95397, which stimulates HIV-1 expression in both J-Lat clonal cell lines, IBET151 only stimulates GFP expression in 10.6 cells (Figure 1G). Through our screen and follow-up validation, we identified NSC95397, a compound not previously linked to HIV-1 latency reversal, which reactivates HIV-1 expression in multiple latency clones with unique integration events.

**Figure 1.**
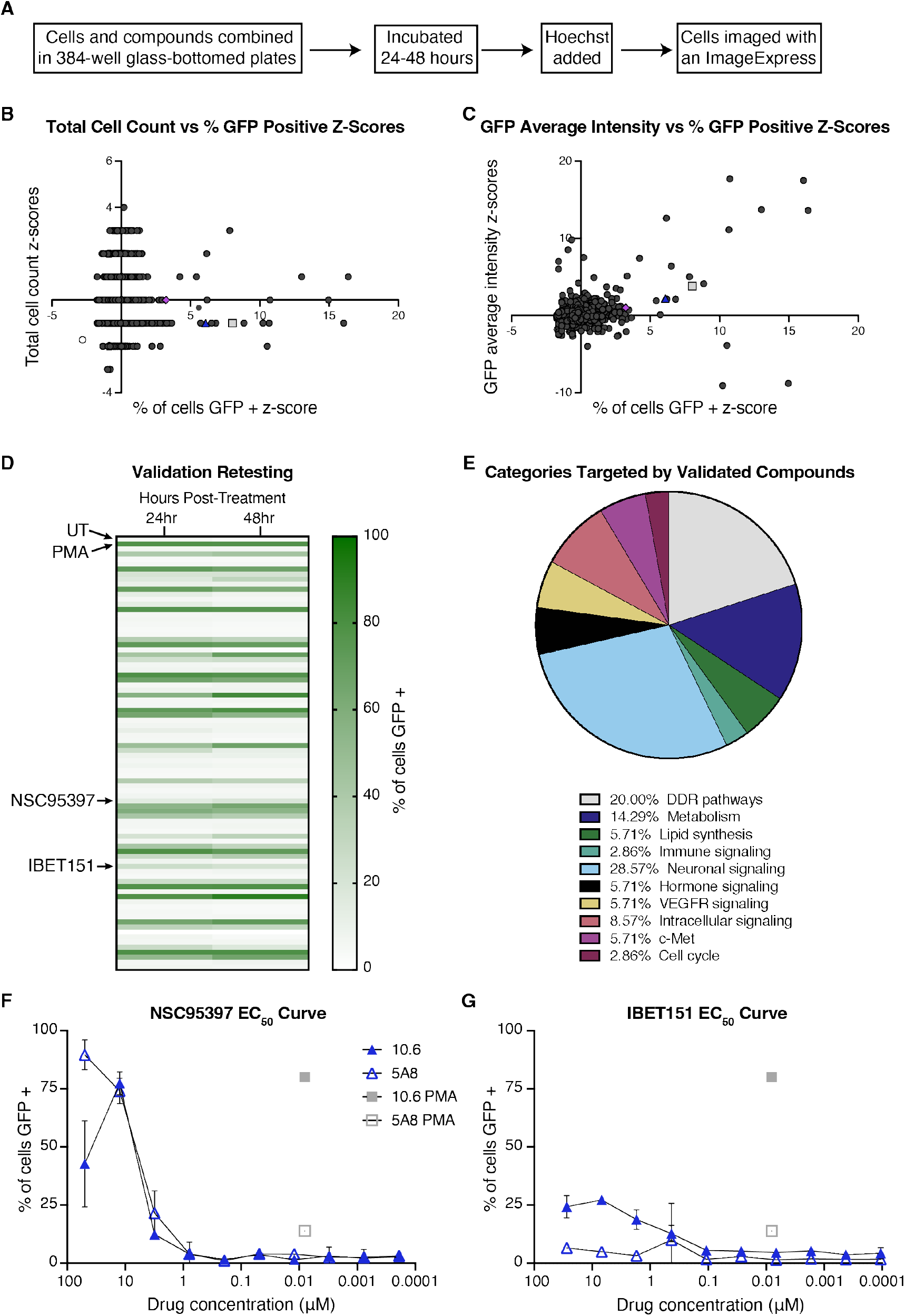
NSC95397 identified as a HIV-1 latency reversal agent (LRA) from a medium-throughput screen. **A)** Schematic detailing the screen methodology. **B-C)** Z-scores of total cell count or GFP intensity vs percent of cells GFP positive. The grey square and purple diamond are two known latency reversal agents, PMA and IBET151, respectively, and the blue triangle is a previously unknown LRA, NSC95397. **D)** 83 compounds were rescreened in triplicate at 24 and 48 hours. Each row of the heatmap represents a different compound, with the top and second row being the controls: untreated (UT) and PMA. **E)** A circle graph representing the pathways/targets of the 35 compounds that revalidated in **D. F-G)** The EC_50_ titration of NSC95397 and IBET151 in J-Lat 10.6 and 5A8 cells. PMA was used as a positive control at 50.0 nM.

### NSC95397 induces both GFP and p24 expression in multiple J-Lat clones

One of the consequences of fluorescence-based screens is the potential for false positives from autofluorescent compounds. To rule out false positives and ensure true HIV-1 reactivation, we assayed hits from our screen for intracellular HIV-1 p24 via flow cytometry, HIV-1 and GFP mRNA expression by qRT-PCR, and GFP protein expression by flow cytometry and western blot. Using flow cytometry, NSC95397 is slightly autofluorescent at higher concentrations in the GFP channel with 30-40% of cells GFP positive (Figure 2A) vs 20% p24 positive (Figure 2B). qRT-PCR confirmed expression of both GFP and unspliced HIV-1 RNA with fold changes ranging 10 – 150 compared to untreated cells (Figure 2 C-D). Lastly, we confirmed protein expression by western blot for GFP in both J-Lat 10.6 and 5A8 cells, further indicating that NSC95397-mediated HIV-1 reactivation occurs irrespective of integration site (Figure 2E). In contrast to NSC95397, another hit from the screen, sunitinib, a receptor tyrosine kinase inhibitor used to treat cancer (22), has high levels of “GFP” via flow cytometry (Figure S2A, S1C-E). However, in those same cells no intracellular HIV-1 p24 is detected (Figure S2B). Consistent with the lack of protein expression, there is no gene expression for GFP or HIV-1 as measured by qRT-PCR (Figure S2D-E). Lastly, autofluorescent (“green/GFP”) cells are identified in sunitinib-treated Jurkat cells by flow cytometry (Figure S2C). These data indicate that while sunitinib is a false positive with autofluorescence in the GFP channel, NSC95397 induces true HIV-1 RNA and protein expression in multiple J-Lat clones.

**Figure 2.**
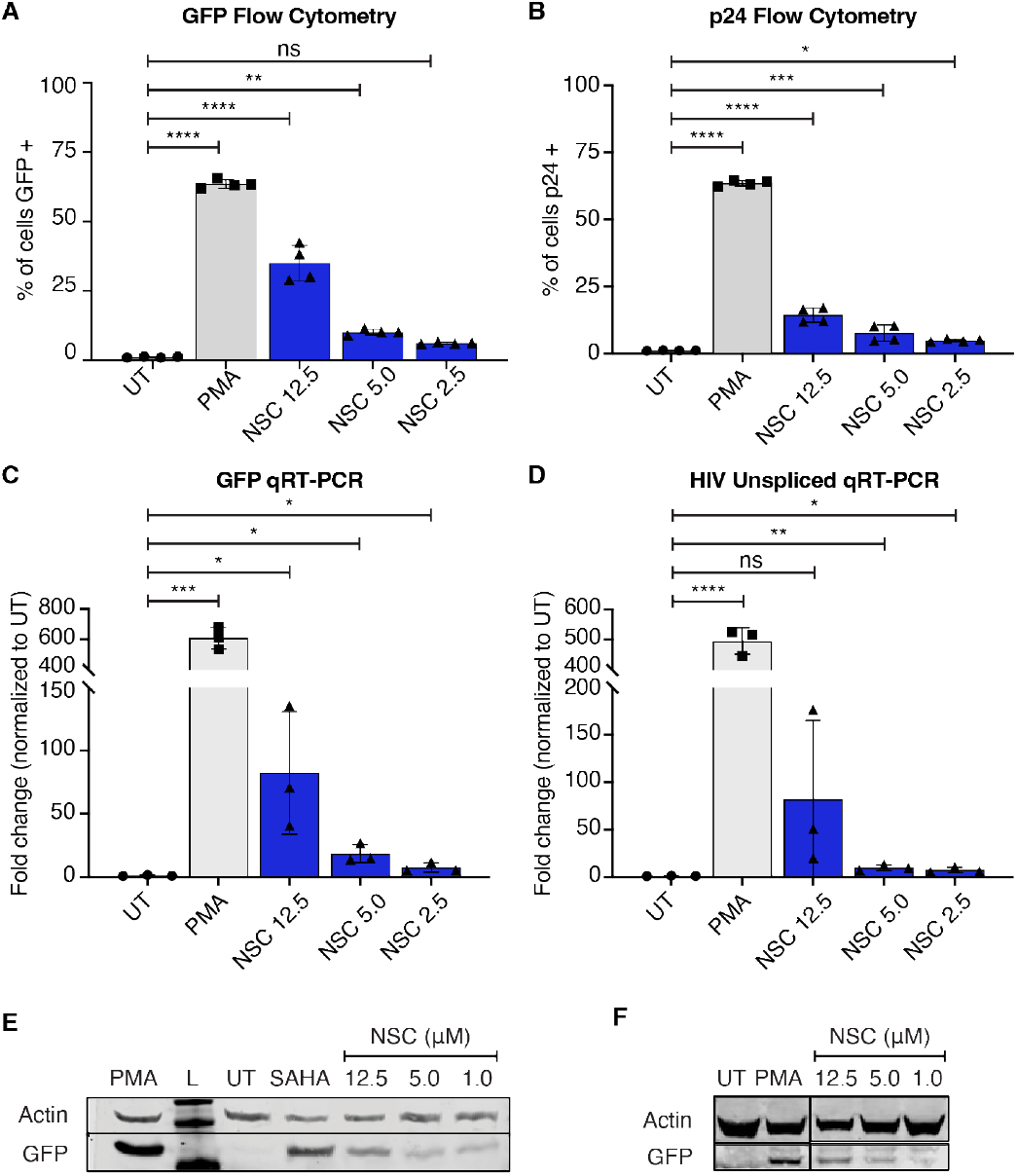
NSC9597 reactivates GFP and p24 expression in J-Lat 10.6 and 5A8 cells. J-Lat 10.6 (**A-E**) or 5A8 (**F**) cells were untreated or treated with 5.0 ng/mL PMA (positive control), or a titration of NSC95397 (NSC) from 12.5 μM to 1.0 μM. **A-B)** Bar graphs showing GFP expression (**A**) or intracellular HIV-1 p24 (**B**) measured via flow cytometry. **C-D)** Bar graphs showing GFP (**C**) or unspliced HIV-1 RNA transcripts (**D**) measured via qRT-PCR. **E-F)** Western blot of whole cell lysates of either J-Lat 10.6 (**E**) or 5A8 cells (**F**) probed for GFP and actin (loading control). 25.0 μM SAHA was used as appositive control. L = ladder; UT = untreated. Results were analyzed with a one-way ANOVA with Turkey’s multiple comparisons tests in **A-B** or an unpaired t test in **C-D**; ns = not significant, * = p ≤ 0.0332, ** = p ≤ 0.0021, *** = p ≤ 0.0002, **** = p ≤ 0.0001.

### NSC95397 synergizes with known LRAs across different latency models

While NSC95397 has been studied as a dual-specificity phos-phatase inhibitor, it has not been correlated with canonical HIV-1 latency reactivation pathways, nor well-tested in immune cells. Since one of our initial goals was to identify compounds that reverse latency in novel mechanisms and work in combination with current LRAs, we first looked to see whether NSC95397 could synergize with other known latency reversing compounds. Compounds that synergize can be used at lower concentrations for a given effect and ideally decrease side effects and toxicity. Synergy can also help to inform molecular mechanisms, as we expect compounds that act in different pathways to have greater potential for synergy when compared to the largely additive effects from compounds that act in overlapping pathways.

We first assessed the potential for NSC95397 to synergize with SAHA, a histone-deacetylase inhibitor. At 12.5 to 1.0 µM, SAHA alone reactivates both J-Lat 10.6 and 5A8 cells in a titratable manner (Figure 3A-B). When combined with 1.5 µM NSC95397 in the J-Lat 10.6 cells, there is a synergistic increase of reactivation with 5.0 µM and 1.0 µM SAHA (Figure 3A, Table 1). In the J-Lat 5A8 cells, there is also synergy at 5.0 µM SAHA (Figure 3B, Table 1).

**Table 1.** Summary of NSC95397 and SAHA combination index synergy calculations. CI (combination index) values for drug-drug effect were calculated using CompuSyn. Concentrations with CI ≤ 0.90 are considered to be synergistic (green)- or greater than additive (with lower values indicating stronger synergy). Concentrations close to 1.0 are considered to have an additive effect (white), in which the two drugs exhibit the same effect from an independent dosage. Concentrations with CI ≥ 1.10 are considered to be antagonistic (purple), or less than additive (with higher values indicating stronger antagonism). Unless otherwise stated, experiments were done in normoxic conditions.

| Cell Line | NSC95397 [uM] | SAHA [uM] | CI | Drug-Drug Effect |
| --- | --- | --- | --- | --- |
| J-Lat 10.6 | 1.5 | 12.5 | 1.22 | Moderate Antagonism |
|  | 1.5 | 5.0 | 0.85 | Slight Synergism |
|  | 1.5 | 1.0 | 0.89 | Slight Synergism |
| J-Lat 5A8 | 1.5 | 12.5 | 1.13 | Slight Antagonism |
|  | 1.5 | 5.0 | 0.87 | Slight Synergism |
|  | 1.5 | 1.0 | 0.91 | Nearly Additive |
| ACH-2 | 2.5 | 10.0 | 0.75 | Moderate Synergism |
|  | 2.5 | 2.0 | 0.50 | Synergism |
|  | 2.5 | 0.5 | 0.88 | Slight Synergism |
| U1 | 2.5 | 10.0 | 1.49 | Antagonism |
|  | 2.5 | 2.0 | 0.18 | Strong Synergism |
|  | 2.5 | 0.5 | 0.10 | Strong Synergism |
| J-Lat 10.6 (Hypoxia) | 1.5 | 12.5 | 0.88 | Slight Synergism |
|  | 1.5 | 5.0 | 0.67 | Synergism |
|  | 1.5 | 1.0 | 0.67 | Synergism |
| J-Lat 5A8 (Hypoxia) | 1.5 | 12.5 | 1.01 | Nearly Additive |
|  | 1.5 | 5.0 | 0.79 | Moderate Synergism |
|  | 1.5 | 1.0 | 1.06 | Nearly Additive |

**Figure 3.**
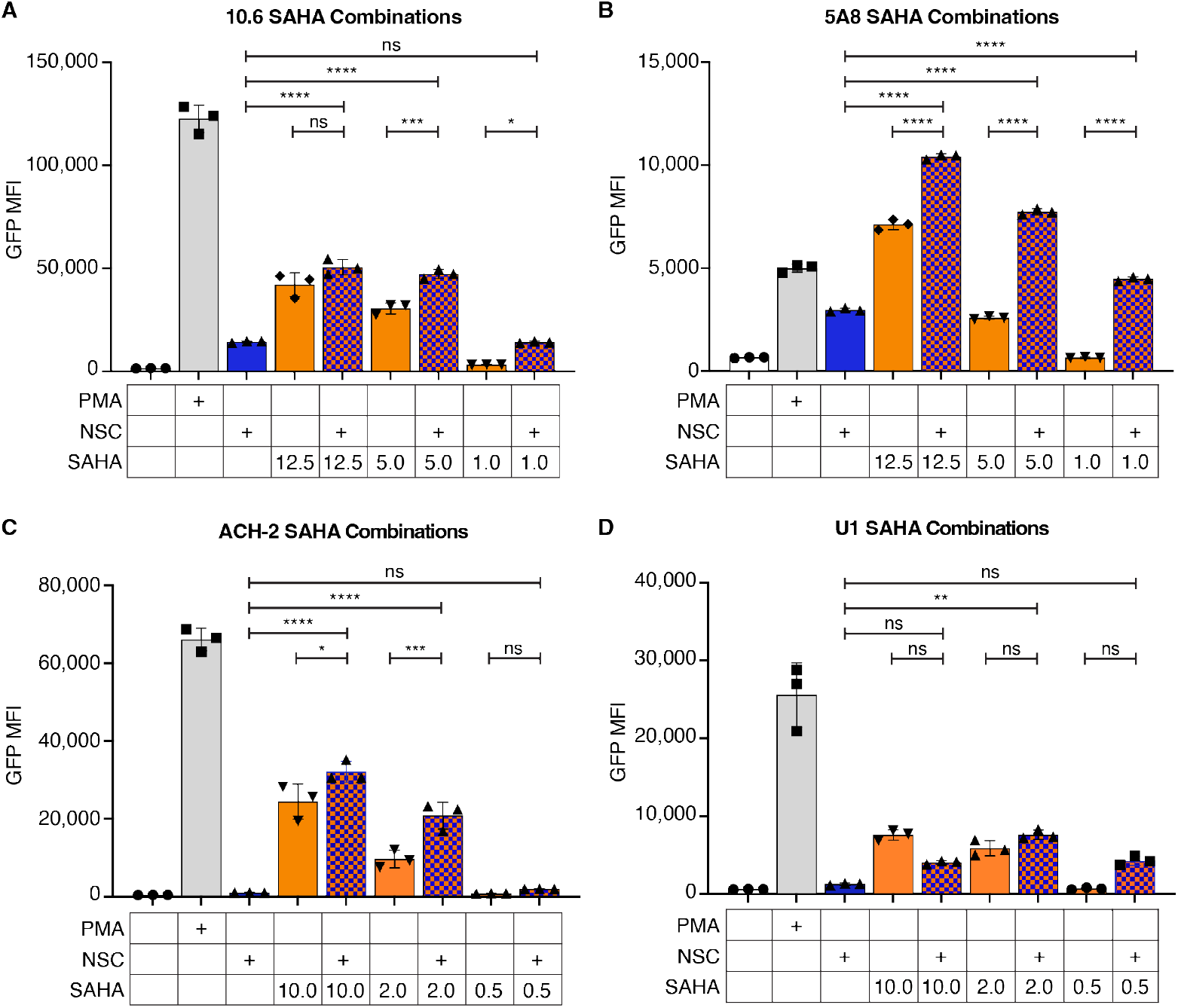
NSC95397 increases HIV-1 reactivation in combination with SAHA. **A-B)** J-Lat 10.6 (**A**) or 5A8 (**B**) cells were untreated or treated with 5.0 ng/mL PMA, 1.5 μM NSC95397 (NSC), a titration of SAHA from 12.5 to 1.0 μM, or a titration of SAHA with NSC held constant at 1.5 μM. **C-D)** ACH-2 (**C**) or U1 (**D**) cells were untreated or treated with 5.0 ng/mL (**C**) or 20 ng/mL PMA (**D**), 2.5 μM NSC, a titration of SAHA from 10 to 0.5 μM, or a titration of SAHA with NSC held constant at 2.5 μM. Bar graphs show GFP mean fluorescence intensity (MFI) measured via flow cytometry. Results were analyzed with a one-way ANOVA with Turkey’s multiple comparisons tests; ns = not significant, * = p ≤ 0.0332, ** = p ≤ 0.0021, *** = p ≤ 0.0002, **** = p ≤ 0.0001.

To determine if this phenotype is maintained across different models of latency, including other important reservoir cell types (such as monocytes) and more diverse (non-clonal) integration sites, we assayed for the ability of NSC95397 to synergistically reactivate HIV-1 from ACH-2 and U1 cell lines. ACH-2 and U1 are persistently infected T and monocytic cell lines, respectively, that are functionally Tat deficient, express low levels of HIV-1 which can be increased by stimulation with LRAs, and have a diversity of non-clonal integration sites (18, 23) making them vastly different models of latency from J-Lat cell lines. At 10.0 to 0.5 µM, SAHA alone reactivates both ACH2 and U1 cells in a titratable manner (Figure 3C-D). Using 2.5 µM NSC95397, and SAHA ranging from 10.0 to 0.5 µM, we see synergistic reactivation at the 2.0 and 0.5 µM in both ACH-2 and U1 cells (Figure 3C-D, Table 1). Of note, we also found that NSC95397 and SAHA together increased the percentage of reactivated dead cells at certain concentrations for all cell lines tested (Figure S3). Together, this suggests that NSC95397 is synergistic with SAHA across multiple diverse models of HIV-1 latency and unlikely to act in the same pathway.

We next assessed the potential for NSC95397 to synergize with prostratin, a protein kinase C (PKC) agonist that reactivates latent HIV-1 through NF-κB mediated transcriptional activation. In J-Lat 10.6 cells, 0.66 to 0.1 µM of prostratin alone leads to a titratable increase of GFP compared to untreated cells (Figure 4A, S4A). Similarly, 5.0 to 0.5 µM of prostratin has a titratable increase of GFP in J-Lat 5A8 cells compared to untreated cells (Figure 4B, S4B). When combining the lower concentrations of prostratin with 1.5 µM NSC95397, there is a synergistic effect for the 10.6 and an additive effect in the 5A8 cells (Figure 4A-B, S4A-B, Table 2).

**Table 2.** Summary of NSC95397 and prostratin combination index synergy calculations. CI (combination index) values for drug-drug effect were calculated using CompuSyn. Concentrations with CI ≤ 0.90 are considered to be synergistic (green), or greater than additive (with lower values indicating stronger synergy). Concentrations close to 1.0 are considered to have an additive effect (white), in which the two drugs exhibit the same effect from an independent dosage. Concentrations with CI ≥ 1.10 are considered to be antagonistic (purple), or less than additive (with higher values indicating stronger antagonism). Unless otherwise stated, experiments were done in normoxic conditions.

| Cell Line | NSC95397 [uM] | Prostratin [uM] | CI | Drug-Drug Effect |
| --- | --- | --- | --- | --- |
| J-Lat 10.6 | 1.5 | 0.660 | 1.00 | Near Additive |
|  | 1.5 | 0.330 | 0.84 | Moderate Synergism |
|  | 1.5 | 0.100 | 0.62 | Synergism |
| J-Lat 5A8 | 1.5 | 5.000 | 2.63 | Antagonism |
|  | 1.5 | 1.000 | 1.05 | Near Additive |
|  | 1.5 | 0.500 | 0.92 | Near Additive |
| ACH-2 | 2.5 | 0.500 | 1.00 | Near Additive |
|  | 2.5 | 0.250 | 0.76 | Moderate Synergism |
|  | 2.5 | 0.125 | 0.65 | Synergism |
| U1 | 2.5 | 5.000 | 2.99 | Antagonism |
|  | 2.5 | 1.000 | 0.44 | Synergism |
|  | 2.5 | 0.500 | 0.30 | Synergism |
| J-Lat 10.6 (Hypoxia) | 1.5 | 0.660 | 1.01 | Near Additive |
|  | 1.5 | 0.330 | 0.84 | Moderate Synergism |
|  | 1.5 | 0.100 | 0.85 | Slight Synergism |
| J-Lat 5A8 (Hypoxia) | 1.5 | 5.000 | 1.07 | Near Additive |
|  | 1.5 | 1.000 | 0.81 | Moderate Synergism |
|  | 1.5 | 0.500 | 0.79 | Moderate Synergism |

**Figure 4.**
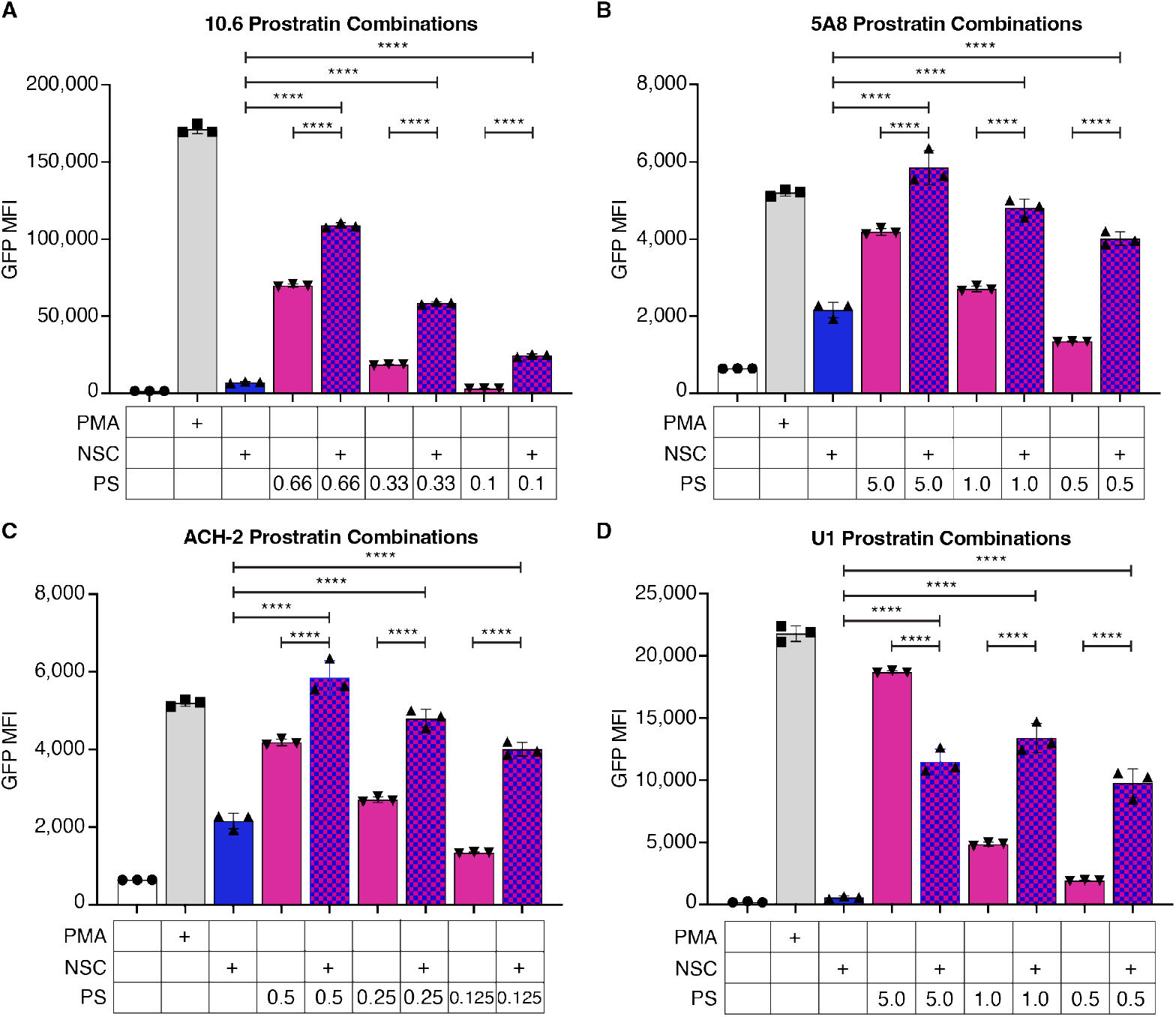
NSC95397 increases HIV-1 reactivation in combination with prostratin. **A-B)** J-Lat 10.6 (**A**) or 5A8 (**B**) cells were untreated or treated with 5.0 ng/mL PMA, NSC95397 (NSC) at 1.5μM, a titration of prostratin (PS) from 0.66 to 0.1 μM (**A**) or 5.0 to 0.5 μM (**B**), or a titration of PS with NSC held constant at 1.5 μM. **C-D)** ACH-2 (**C**) or U1 (**D**) cells were untreated or treated with 5.0 ng/mL (**C**) or 20.0 ng/mL PMA (**D**), 2.5 μM NSC, a titration of PS from 0.5 to 0.125 μM (**C**) or 5.0 to 0.5 μM (**D**), or a titration of PS with NSC held constant at 2.5 μM. Bar graphs show GFP MFI measured via flow cytometry. Results were analyzed with a one-way ANOVA with Turkey’s multiple comparisons tests; ns = not significant, * = p ≤ 0.0332, ** = p ≤ 0.0021, *** = p ≤ 0.0002, **** = p ≤ 0.0001.

We repeated this in ACH-2 cells with 2.5 µM NSC95397 and prostratin ranging from 0.5 to 0.125 µM, as well as U1 cells with 2.5 µM NSC95397 and prostratin ranging from 5.0 to 0.5 µM (Figure 4C-D). As with the J-Lat 10.6 cells, we see synergistic reactivation at the lower two concentrations for both ACH-2 and U1 cells (Figure 4C-D, Table 2). Similar to SAHA, NSC95397 in combination with prostratin leads to a slight increase of reactivated dead (Figure S4). Together, this suggests that prostratin and NSC95397 act in different, but potentially synergistic, pathways irrespective of integration site and cell type.

### NSC95397 does not increase global histone markers of open chromatin

The evidence of synergy with NSC95397 and SAHA suggests, that unlike SAHA, NSC95397 does not reactivate latent HIV-1 by modifying the global epigenetic landscape to increase euchromatin. To directly test this, we treated J-Lat 10.6 and 5A8 cells with NSC95397 and assessed the levels of three markers of active promoter and enhancer regions by flow cytometry: H3K4me3, H3K9ac, H3K27ac. As expected, in both cell lines, there is no change for all three markers of active chromatin in NSC95397 treated cells when compared to untreated and PMA-treated cells (Figure 5, S5). This is opposed to SAHA-treated cells, which significantly increased H3K4me3, H3K9ac, and H3K27ac. This further supports the hypothesis that NSC95397 does not act in a manner similar to SAHA to alter the global chromatin landscape to euchromatin.

**Figure 5.**
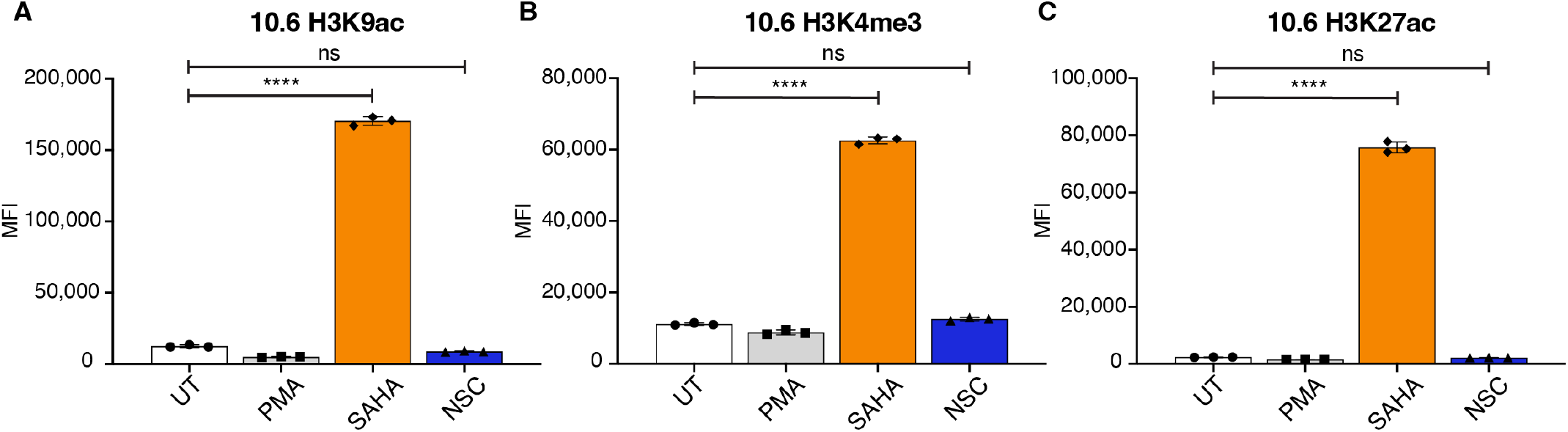
NSC95397 does not increase global histone markers of open chromatin. **A-C)** J-Lat 10.6 cells were untreated or treated with 5.0 ng/mL PMA, 25.0 μM SAHA (positive control), or NSC95397 (NSC) at 2.5 μM. Bar graphs show H3K9ac (**A**), H3K4me3 (**B**), or H3K27ac (**C**) MFI measured by flow cytometry. Bar graphs show GFP MFI measured via flow cytometry. Results were analyzed with a one-way ANOVA with Turkey’s multiple comparisons tests; ns = not significant, * = p ≤ 0.0332, ** = p ≤ 0.0021, *** = p ≤ 0.0002, **** = p ≤ 0.0001.

### NSC95397 does not increase known HIV-1-associated transcription pathways

As NSC95397 does not appear to change histone modifications nor act through PKC/NF-κB transcription, we utilized bulk RNA sequencing to identify potential transcriptional pathways associated with NSC95397-mediated latency reversal. J-Lat 10.6 cells were treated with 5.0 µM NSC95397 for 24 hours, with 5.0 ng/mL PMA and 25.0 µM SAHA as positive controls. Consistent with their different effects on the cell, PMA, SAHA, and NSC95397 treated cells group independently in both PCA and cluster analysis (Figure 6A-B). Moreover, NSC95397 is indistinguishable from untreated control cells (Figure 6A-B), suggesting NSC95397 does not alter the global transcriptome. PMA, SAHA, and NSC95397 all increase viral transcripts compared to untreated cells (Figure 6C, S6A, S7A-B).

**Figure 6.**
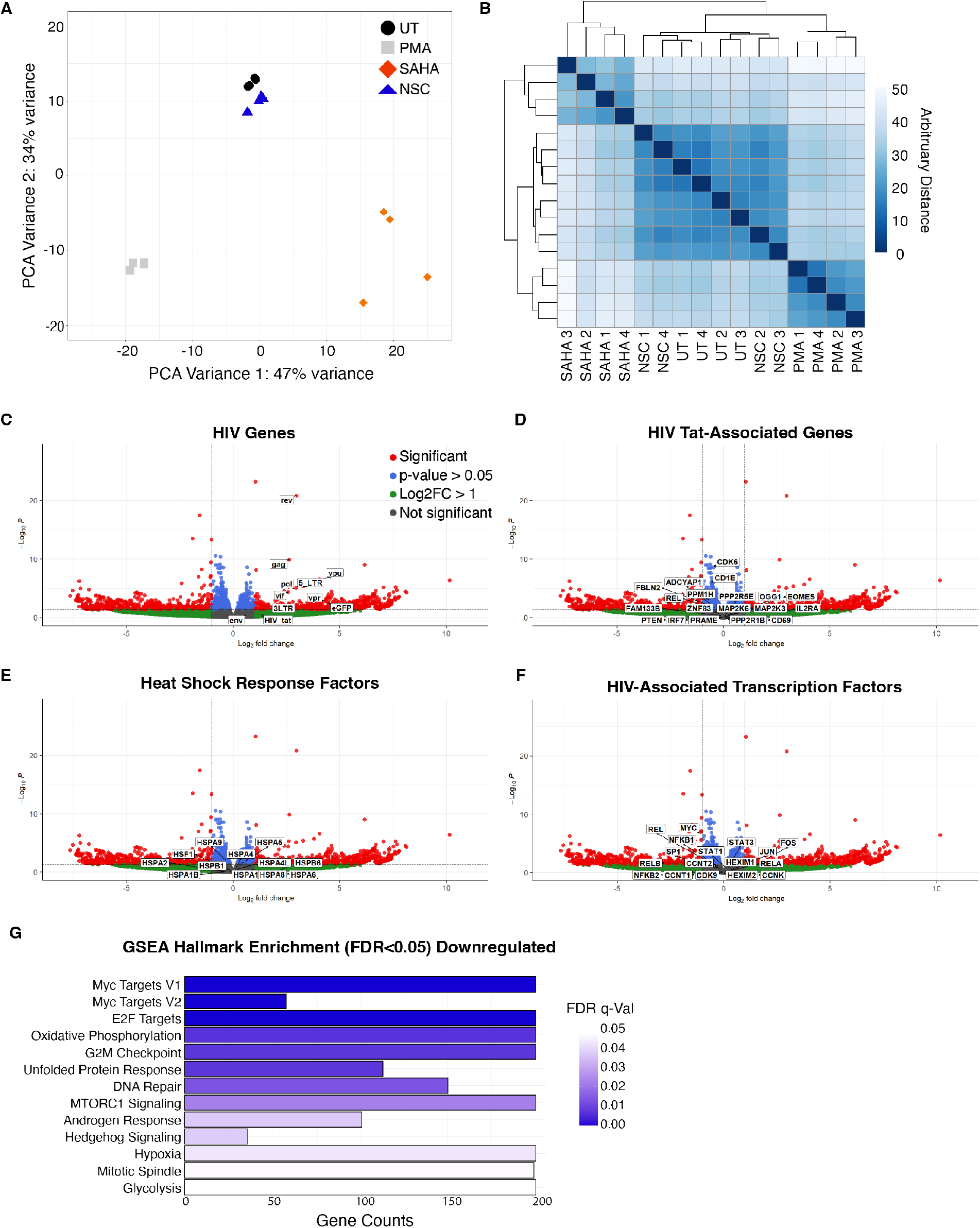
NSC95397 does not globally activate HIV-1 latency reversal associated pathways. J-Lat 10.6 cells were untreated or treated with 5.0 ng/mL PMA, 25.0 μM SAHA, or 5.0 μM NSC95397 (NSC) and analyzed by bulk RNA sequencing in four biological replicates. **A)** Principal component analysis (PCA) of all normalized transcripts in all samples. **B)** Clustered heatmap of all normalized transcripts in all samples. **C-F)** Volcano plot labeling HIV-associated reads (**C**), selected HIV-1 Tat-modulated genes (**D**), key heat shock response genes (**E**), and HIV-associated transcription factors (**F**). Cut-offs are drawn for log2 fold change above 1 and p-value greater than 0.05 where reads are separated to non-significant and non-enriched (gray), non-significant with log2 fold change above 1.0 (green), significant and log2 fold change below 1.0 (blue), or significant and log2 fold change above 1.0 (red). **G)** Gene ontology using Gene Set Enrichment Analysis (GSEA) identifying top negatively enriched hallmarks.

Given the lack of global changes in the transcriptome, we assessed whether NSC95397 alters HIV transcription through other known mechanisms. As HIV-1 Tat stimulates LTR transcription and elongation, we also looked for upregulation in Tat-regulated host genes but found little change and correlation to Tat-mediated host transcripts (Figure 6D), consistent with the results in ACH-2 and U1 cells, confirming NSC95397 does not specifically work through stimulation of HIV-1 Tat. As some compounds may reactivate latently infected cells due to heat shock stress (24), we also looked for increase in heat shock response transcripts but saw little change in NSC95397 treated cells (Figure 6E).

Notably, unlike the positive controls, NSC95397 had few differentially expressed genes (DEGs) outside of the HIV-1 transcripts, suggesting that NSC95967 may specifically activate the latent HIV-1 and few other genes. Gene Set Enrichment Analysis (GSEA) for enriched hallmarks did not identify any positively enriched hallmarks for NSC95397. This is supported by the lack of increased HIV-related transcription factor transcripts (Figure 6F) as well as the lack of enrichment of transcription factors by TRRUST (transcriptional regulatory relationships unraveled by sentence-based text-mining, http://www.grnpedia.org/trrust) analysis (Figure S6B). While TRRUST also did not show enrichment of transcription-factor-mediated changes for downregulated genes and pathways (Figure S6C), GSEA analysis identified significant negatively enriched hallmarks, including metabolism, cell cycle, hypoxia/redox, stress response, and DNA damage/repair (Figure 6G, Supplement File 1). Thus, our RNA sequencing analyses indicate that NSC95397 does not greatly alter the global transcriptional landscape, despite specifically activating HIV-1 transcription..

### NSC95397 increases markers of DNA damage

DNA damage has been linked to transcriptional and epigenomic changes, some of which have been correlated with latency reversal (25, 26). Since NSC95397 stimulation downregulates genes in pathways mediating DNA repair (Figure 6G), we asked if NSC95397 treatment leads to an accumulation of DNA damage. We treated U2OS cells with DMSO (vehicle control), Etoposide (positive control for DNA breaks), PMA, or NSC95397 then probed for three different markers of the DNA damage response (DDR) by immunofluorescence: γH2AX (general DNA damage), RPA32 (single-stranded DNA lesions), and 53BP1 (double-stranded DNA breaks). NSC95397 treated cells have an increase in γH2AX mean fluorescence intensity when compared to untreated cells, though to a much lesser extent than Etoposide (Figure 7). Similar results are seen for RPA32 and 53BP1 when measuring foci per cell; however, the effect on 53BP1 is not as robust as RPA32, suggesting potential specificity in the DDR activation by NSC95397 (Figure 7). Together, this supports our RNA sequencing analysis and suggests that NSC95397 activates the host DDR, particularly single-stranded DNA lesion pathways.

**Figure 7.**
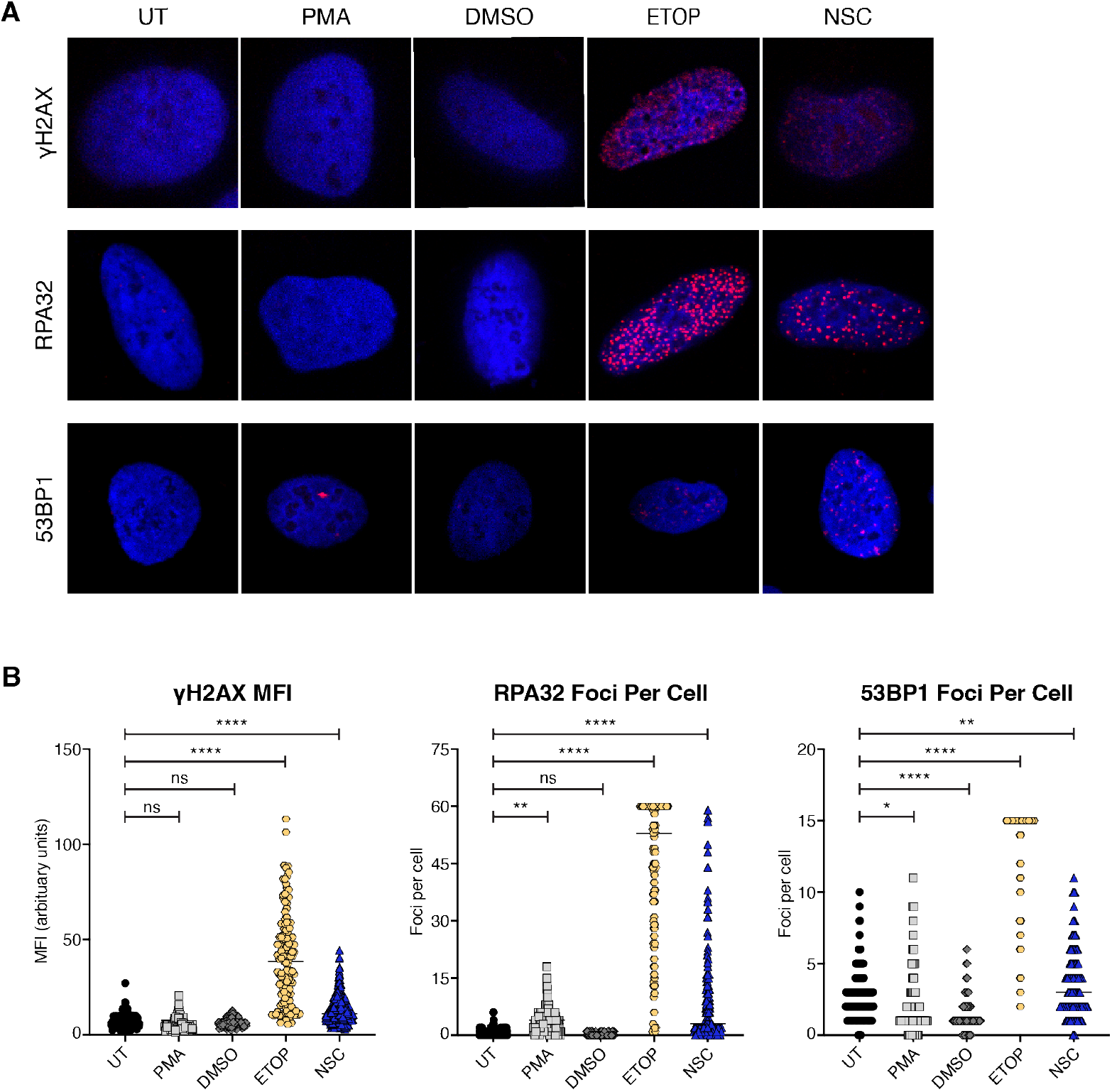
NSC95397 causes accumulation of DNA damage markers. **A)** Representative images of U2OS cells untreated or treated with 10.0 μL DMSO (negative control), 50.0 μM Etoposide (ETOP; DNA damage positive control), 5.0 ng/mL PMA, or 5.0 μM NSC95397 (NSC) for 24 hours and assessed for presence of DNA damage markers by confocal microscopy. Blue (Hoechst 33342) shows nuclei while damage markers (γH2AX, RPA32, or p53BP1) are in red. **B)** Mean fluorescence intensity (MFI) of at least 100 cells per condition was quantified for γH2AX. Foci per cell of at least 100 cells per condition was quantified for RPA32 and 53BP1 where foci above the ETOP control are binned at 60 (RPA32) or 15 (53BP1). Results were analyzed with a one-way ANOVA with Dunnett multiple comparisons tests; ns = not significant, * = p ≤ 0.0332, ** = p ≤ 0.0021, *** = p ≤ 0.0002, **** = p ≤ 0.0001.

### NSC95397 increases HIV-1 reactivation in combination with established LRAs in hypoxic conditions

A major difficulty in identifying viable LRAs is that initial studies are often done in cells cultured at non-physiological conditions such as high oxygen (normoxia). This results in large discrepancies when potential therapeutic LRAs are tested at more physiological levels of low oxygen (hypoxia) that more closely recapitulate the physiological conditions in tissues that harbor latent cells and in clinical trials (20). Given this importance, and that our RNA sequencing result identified a downregulation of response to hypoxia and other metabolic pathways (Figure 4G), we asked if NSC95397 reactivates latent HIV-1 when cells are maintained in hypoxic conditions. J-Lat 10.6 and 5A8 cells were maintained at either normoxic (21% O_2_) or hypoxic (1% O_2_) culture conditions while treated with LRAs for 24 hours. While reactivation with PMA in both J-Lat 10.6 and 5A8 clonal lines decreases in hypoxia, NSC95397 activates both J-Lat 10.6 and 5A8 cells in a similar titratable manner between normoxic and hypoxic conditions (Figure 8A-B, and S8A-B), with only high concentrations of NSC953937 in 5A8 cells showing statistical differences between the two conditions (Figure 8B, S8B). Similar to normoxic conditions (Figure S3), experimental groups treated with NSC95397 also show an increase of the reactivated dead population in both cell lines when maintained under hypoxic conditions (Figure S8).

**Figure 8.**
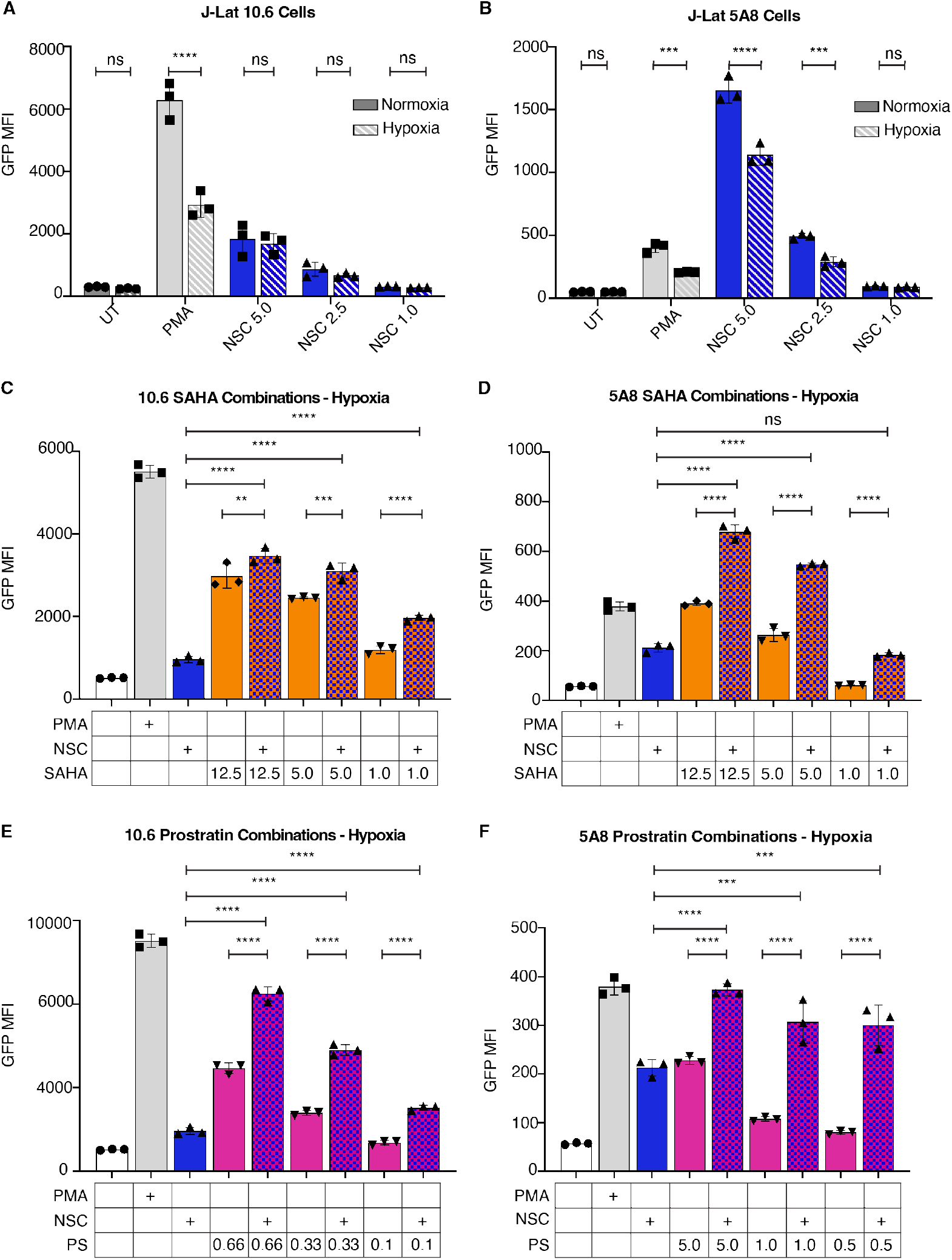
NSC95397 increases HIV-1 reactivation in hypoxic conditions. **A-B)** Bar graphs showing GFP Mean Fluorescence Intensity (MFI) for J-Lat 10.6 (**A**) or 5A8 (**B**) cells. Cells were untreated, treated with 5.0 ng/mL PMA (positive control), or a titration of NSC95397 (NSC) from 5.0 μM to 1.0 μM. Solid bars are cells maintained in normoxia while patterned bars are cells maintained in hypoxia. **C-D)** J-Lat 10.6 (**C**) or 5A8 (**D**) cells were grown in hypoxic conditions and untreated or treated with 5.0 ng/mL PMA (positive control), 1.5 μM NSC, or NSC held constant at 1.5 μM with a titration of SAHA from 12.5 to 1.0 μM. Bar graphs show GFP mean fluorescence intensity (MFI) measured via flow cytometry. **E-F)** J-Lat 10.6 (**E**) or 5A8 (**F**) cells were untreated or treated with 5.0 ng/mL PMA, NSC at 1.5 μM, or NSC held constant at 1.5 μM with a titration of prostratin (PS) from 0.66 to 0.1 μM (**E**) or 5.0 to 0.5 μM (**F**). Bar graphs show GFP MFI measured via flow cytometry. Results were analyzed with a two-way ANOVA with Šídák’s multiple comparisons test; ns = not significant, * = p ≤ 0.0332, ** = p ≤ 0.0021, *** = p ≤ 0.0002, **** = p ≤ 0.0001.

We next tested whether combination of NSC95397 with SAHA or prostratin further increased reactivation under hypoxia, similar to normoxia (Figure 3 and 4). Consistent with normoxia conditions, there is a synergistic increase of reactivation when NSC95397 is combined with SAHA in both J-Lat 10.6 and 5A8 cells (Figure 8C-D, S8C-D, Table 1), though the absolute levels are not as high as normoxia. When NSC95397 is combined with prostratin, there is a synergistic increase of reactivation in both J-Lat 10.6 and 5A8 cells (Figure 8E-F, S8 E-F, Table 2). Together, our data suggests that NSC95397 alone and in combination robustly reactivates cells under more physiological hypoxic settings.

## Discussion

In this study, we identified NSC95397 as a novel LRA from a screen of over 4000 compounds in J-Lat 10.6 cells. We show that NSC95397 activates HIV-1 gene expression in multiple J-Lat T cell clones, polyclonal ACH-2 T cells, and polyclonal U1 premonocytic cells. In all four cell lines, NCS95397 synergized with known LRAs, SAHA and prostratin, suggesting it reactivates latent HIV-1 through distinct mechanisms from these drugs. In line with this, NSC95397 did not globally alter histone markers of open chromatin or the transcriptome, despite HIV-1 reactivation. However, NSC95397 negatively enriched hallmarks for metabolism, cell cycle, hypoxia/redox, stress response, and DNA damage/repair, the latter of which was validated by immunofluorescence for DDR activation markers. Finally, NSC95397 synergized with both SAHA and prostratin in physiologically relevant hypoxic conditions. Together, our data suggests NSC95397 is a novel LRA that exerts minimal effects on cells and acts through unknown mechanisms.

As an LRA, NSC95397 has unique and promising characteristics for use in HIV-1 cure strategies. Unlike established LRAs such as PMA and SAHA, NSC95397 does not induce widespread transcriptional activation, which could limit off-target effects and side effects, making it a potentially safer option for therapeutic use. Our RNA sequencing data indicate that at the concentrations and timing used, NSC95397 causes very few differentially expressed genes, leaving the overall transcriptome largely indistinguishable from untreated cells. Additionally, the lack of changes in global histone markers of open chromatin supports the idea that NSC95397 exerts minimal effects on global transcription and chromatin landscape, further suggesting it may cause fewer side effects compared to other LRAs.

Some studies have demonstrated that LRAs may have difficulty stimulating HIV-1 reactivation *in vivo* due in part to differential transcription and epigenetic control of diverse integration sites (5, 27). This challenge is further compounded by the diversity of reservoir cell types, integration sites, and the tissue microenvironment (20, 28, 29). However, NSC95397 shows promise in overcoming these barriers, particularly due to its ability to reactivate HIV-1 latency in cells maintained under hypoxic conditions, which is a more physiologically relevant environment (Figure 8). Additionally, our findings revealed that NSC95397 synergizes with current LRAs, such as SAHA and prostratin, even at low concentrations (Figures 3, S3, 4, S4, and 8). This synergy allows for the use of lower concentrations of each compound in combination therapies, maximizing efficacy while minimizing toxicity and other side effects. Low concentrations of NSC95397 in combination with low concentrations of SAHA and prostratin is synergistic in all models used, as well as in the more physiologically relevant hypoxic environment. Interestingly, the two J-Lat clonal cell lines and the U1 and ACH-2 pooled lines we used respond, both in HIV-1 expression and cell toxicity, to NSC95397 co-treatments differentially (Figure S3, S4, and S7). The differential responses observed across various cell models to NSC95397 co-treatments highlight the issue of reservoir heterogeneity. This underscores the importance of developing LRA cocktails that target multiple pathways to effectively reach a greater proportion of the latent viral reservoir. Given these characteristics, NSC95397 or future derivatives could be valuable additions to combination therapy strategies, addressing the complexities posed by reservoir diversity while enhancing overall treatment effectiveness.

While NSC95397 shows potential as a cure therapy, the exact mechanisms by which it reactivates latent HIV-1 remain to be fully elucidated. As a broad dual-specificity phosphatase inhibitor, NSC95397 likely influences multiple signaling pathways. Our findings suggest that NSC95397 does not activate transcription pathways typically associated with HIV-1 latency reversal, with the exception of slight downregulation of Myc signaling (2, 3, 30). However, its ability to reverse latency without global transcriptional changes suggests that NSC95397 may act through mechanisms either specific to the LTR or not yet associated with latency modulation. If NSC95397 reactivates latent HIV-1 through LTR-specific mechanisms, targeted studies of the HIV-1 LTR will be necessary to identify the transcription factors and chromatin modifications specifically regulated at the LTR by NSC95397. In line with its role as a phosphatase inhibitor, further exploration of changes in global protein abundance, phosphorylation, and other post-translational modifications may reveal the protein networks affected by NSC95397. Understanding these networks could elucidate how NSC95397 reverses HIV-1 latency and potentially uncover new pathways involved in HIV-1 latency and reactivation.

The repression of pathways related to metabolism, DNA repair, and hypoxia/stress response observed in our RNA sequencing data indicates that changes in the metabolic microenvironment may also play a role in latency modulation. Of particular interest is the downregulation of DNA repair pathways, given the connection between the DDR and HIV-1 replication. Early studies with UV have shown that DNA damage can increase HIV-1 transcription (31, 32). Similarly, members of the DDR have been both directly and indirectly implicated in mechanisms related to latency modulation (26, 33, 34). The inhibition of the DDR by NSC95397 could lead to the accumulation of DNA damage markers and enhanced transcription from viral proteins, contributing to the reactivation of latent HIV-1. The increased cell death observed following reactivation suggests that NSC95397 may sensitize cells to DNA damage post-reactivation, further supporting the need for studies to clarify the role of the DDR in latency reversal.

As the population of PLWH ages, the incidence of both AIDS-related and non-AIDS-related cancers is rising, with cancer now being a leading cause of death among HIV-1 positive individuals on ART (35-37). The interaction between chemotherapy and the latent HIV-1 reservoir remains poorly understood, underscoring the importance of studying how cancer therapies impact HIV-1 infection and latency. Although NSC95397 is not currently used as a cancer treatment, its effects on HIV-1 latency reactivation in our study suggest it could be an interesting candidate for further testing in HIV-1 infected individuals who develop cancer. Additionally, our findings lay the groundwork for evaluating other cancer therapies for their impact on the latent HIV-1 reservoir, potentially opening new avenues for co-therapy in HIV-1 and cancer.

In summary, our results introduce NSC95397 as a novel LRA with significant potential for HIV-1 cure strategies due to its ability to synergize with existing LRAs and its minimal impact on global transcription and chromatin landscapes. However, further studies are needed to fully understand the specific mechanisms by which NSC95397 reactivates latent HIV-1, particularly its effects on the DNA damage response and metabolism. Additionally, future research should explore the effects of NSC95397 on both infected and uninfected cells and evaluate its efficacy as an LRA *in vivo*. These investigations will be crucial for advancing our understanding of latency modulation and guiding the development of new LRA-based therapies.

## Acknowledgements

Lorem ipsum dolor sit amet, consectetuer adipiscing elit. Maecenas porttitor congue massa. Fusce posuere, magna sed pulvinar ultricies, purus lectus malesuada libero, sit amet commodo magna eros quis urna. Nunc viverra imperdiet enim. Fusce est. Vivamus a tellus. Pellentesque habitant morbi tristique senectus et netus et malesuada fames ac turpis egestas. Proin pharetra nonummy pede. This paper was typeset with the bioRxiv word template by @Chrelli: www.github.com/chrelli/bioRxiv-word-template

## Author contributions

We would like to thank the UCLA Molecular Screening Shared Resource, especially Bobby Tofig and Constance Yuen, for their help with performing the drug screen and analysis. Flow cytometry was performed in the UCLA Jonsson Comprehensive Cancer Center (JCCC) and Center for AIDS Research Flow Cytometry Core Facility. We thank the UCLA Technology Center for Genomics & Bioinformatics for performing the RNA sequencing library preparation and sequencing. We would also like to thank Jeff Long and Steve Jacobsen (UCLA) for the use of the LSM 980. We thank Warner Greene (the Gladstone Institutes) and Anjie Zhen (UCLA) for generously providing cells used in this study. This research was funded by NIH NIAID grants R01AI147837 (O.I.F.), U54 AI170660-01 (H.E.T.), and a UCLA DGSOM I3T Discovery Seed Grant (O.I.F), with support from the Ruth L. Kirschstein National Research Service Award AI007323 (R.N.D, V.Y.). This work was also supported by the James B. Pendleton Charitable Trust and the McCarthy Family Foundation. The funders had no role in study design, data collection and interpretation, or the decision to submit the work for publication.

## Author Contributions

Experimental Concept and Design: R.N.D., V.Y., R.D., H.E.T., O.I.F.

Experiment Execution: R.N.D., V. Y., Y.K.

Manuscript Writing and Editing: R.N.D., V.Y., O.I.F.

## Materials and Methods

### Cell Lines and Culture

The ACH-2, U1 (both from BEI Resources and generously provided by Dr. Anjie Zhen, UCLA), and J-Lat 10.6 (BEI Resources) and 5A8 (generously provided by Dr. Warner Greene, the Gladstone Institutes) cells were cultured in flasks (Thermo Fisher Nunc™ EasYFlask™) in RPMI 1640 growth medium (L-glutamine, Gibco) with 10% fetal bovine serum (Peak Serum) and 1% penicillin-streptomycin (Gibco). The human bone osteosarcoma epithelial (U2OS) cells were cultured as adherent cells directly on tissue culture plastic (Greiner) in DMEM growth medium (high glucose, L-glutamine, no sodium pyruvate; Gibco) with 10% fetal bovine serum (Peak Serum) and 1% penicillin-streptomycin (Gibco) at 37°C and 5% CO_2_. U2OS cells were harvested using 0.05% Trypsin-EDTA (Gibco).

### Medium-throughput compound screen

J-Lat 10.6 or 5A8 cells were seeded into 384-well black, glass-bottom plates (Greiner) using an Agilent BioTek Multiflo. All compounds were stored at 10 mM in DMSO, and 0.25 µl was added to each well using a Biomek Fx (Beckman Coulter) equipped with a custom pin tool (V&P). Media containing 0.25 µl DMSO or 5.0 ng/mL PMA was added to the first and last two columns, respectively. Cells were incubated for 24, 36, or 48 hours. Cells incubated with Hoechst (1:2000) (Invitrogen H3570) for 15 minutes then each plate was imaged using ImageXpress Micro Confocal High Content Imaging System (Molecular Devices) at 10x magnification. Images were analyzed using the MetaXpress Analysis software, and the raw data was imported into the Collaborative Drug Discovery platform and z-scores were calculated for each compound using standard deviations on each plate separately. The EC_50_ was conducted similarly; each compound was purchased new, resuspended at 10.0 mM in DMSO, then a 10-step 4-fold dilution was performed and applied to cells as above. Datafitting was performed using a 4-parametric Boltzman equation and EC_50_ using standard settings on CDD or Prism. Follow up studies were done manually with IBET151 (Cayman Chemistry 11181), NSC95397 (Cayman Chemistry 21431), PMA (VWR 102515-692), prostratin (Cayman Chemistry 10272), SAHA (Cayman Chemistry 10009929), and sunitinib (Cayman Chemistry 13159).

### Flow cytometry

J-Lat 10.6 or 5A8, ACH-2, or U1 cells were plated in U-bottom 96-well microplates (Corning Inc) at 1×10^6^ cells/mL in plain or drug media for 24 hours. Cells were then incubated with Ghost Dye (Ghost Dye™ Red 780) (1:4000 µL PBS) for 30 min at 4°C, then fixed with 4% PFA for 15 min. Alternatively, cells were fixed with 4% PFA for 10 min, then permeabilized with 0.1% Saponin for 5 minutes and probed with anti-H3K4me3 (Cell Signaling 12064S, 1:150 in 0.1% Saponin), anti-H3K9ac (Cell Signaling C5B11, 1:150 in 0.1% Saponin), anti-H3K27ac (Cell Signaling D5E4, 1:150 in 0.1% Saponin), or anti-p24 (Beckman Coulter 6604667, 1:250 in 0.1% Saponin) for 1 hour, in the dark as necessary. Cells were analyzed with an Attune NxT Flow Cytometer (Thermo Fisher Scientific) and data analysis was performed with FlowJo (BD).

### Hypoxia Assays

J-Lat 5A8 and 10.6 cells were cultured in RPMI 1640 (Corning 10-040-CV) supplemented with 10% FBS (Cytiva SH30071.03) and 1% penicillin/streptomycin (Gibco 15140-122) at 37°C, 21% O2 and 5% CO2. For the latency reversal assays, cells were plated at a density of 0.5 × 10^6^ cells/mL/well in duplicate sets of 24 well plates. Cells were then treated with latency reversal agents, NSC95397 (1.5 µM), PMA (10.0 ng/mL), SAHA (12.5, 5.0, and 1.0 µM), prostratin (5.0, 1.0, and 0.5 µM for J-Lat 5A8; 0.66, 0.33, and 0.1 µM for J-Lat 10.6 cells, Tocris Bioscience 5739) or their respective combinations in the synergy experiments and each treatment was done in triplicates. One set up was maintained at standard normoxic culture conditions (21% O_2_) while the other was maintained under hypoxic conditions (1% O_2_) in a specialized hypoxic workstation, Whitley H35 HEPA Hypoxystation (37°C, 1% O_2_, 5% CO_2_ and 94% N_2_) for 24 hours. At the 24-hour time point, cells were harvested, pelleted, and resuspended in SYTOX AADvanced dead cell stain (1.0 µM; Invitrogen S10274) on ice. Stained cells were immediately analyzed for viability and GFP expression by flow cytometry collected on a BD LSR Fortessa and data analysis was performed with FlowJo (BD).

### Drug Synergy Calculations

Statistical analysis for drug-drug interactions was calculated using CompuSyn (38, 39) following the Chou-Talalay principles for non-constant drug ratios. Corrected values “µCorr” was calculated as in (40) across experimental triplicates, and percent reactivation was inputted as “effect”. Drug combinations were assigned on a scale of anergy to synergy following Combination Index (CI) calculations as in (38) using the scale from the CompuSyn manual.

### Western Blots

J-Lat 10.6 or 5A8 cells were plated in 6-well tissue culture-treated plates (Greiner Bio-one) at (5×10^6^ cells/mL) in plain or drug media for 24 hrs. Cells were then lysed in radioimmunoprecipitation assay (RIPA) buffer (150 mM NaCl, 1% NP-40, 0.5% DOC, 0.1% SDS, 50 mM Tris) for 15 min on ice and clarified by centrifugation at 14,800 x g for 15 minutes. Lysates were boiled in 4x sample buffer (250 mM Tris-HCL, 8% SDS, 0.2% Bromophenol Blue, 40% Glycerol, 20% β-mercaptoethanol) in preparation for SDS-PAGE using 12% Bis-Tris polyacrylamide gels (Invitrogen) and transferred to polyvinylidene difluoride membranes. Immunoblotting was performed using mouse anti-eGFP (Takara 632380, 1:2000), rabbit anti-Actin (Bethyl A300491A, 1:6000), mouse anti-gag (NIH-ARP#3537, 1:1000) followed by donkey anti-rabbit IgG 800CW (IRDye®,1:20000) or donkey anti-mouse IgG 680RD (IRDye®,1:20000) and imaged on a LI-COR Odyssey M.

### RNA Isolation

Total RNA was isolated from 4×10^6^ J-Lat 10.6 cells according to the W.M. Keck Foundation Biotechnology Resource Laboratory at Yale University TRIzol RNA Isolation protocol (41).

### qRT-PCR

250 ng of input RNA was used as template for reverse transcription by SuperScript IV First-Stand Synthesis System (Invitrogen) with Oligo(dT) primers. qRT-PCR was performed with PowerTrack SYBR Green Master Mix (Thermo Fisher Scientific) on the LightCycler 480 System (Roche) with the following primers: (5’ to 3’) HIV-1 unspliced RNA GCGACGAAGACCTCCTCAG and GAGGTGGGTTGCTTTGATAGAGA and GAPDH CAAGATCATCAGCAATGCCT and AGGGATGATGTTCTGGAGAG. Initial enzyme activation at 95°C for 2 minutes, then 40 cycles of 95°C for 15 seconds and 60°C for 60 seconds. mRNA levels were quantified by calculating ΔΔCt. Target transcript Ct values were normalized to the Ct value of the housekeeping gene GAPDH followed by calculating fold change to untreated cells.

### RNA Library Prep and Sequencing

Libraries for RNA-Seq were prepared with KAPA Stranded mRNA-Seq Kit (Cat.KK8420). The workflow consists of mRNA enrichment and fragmentation, first strand cDNA synthesis using random priming followed by second strand synthesis converting cDNA:RNA hybrid to double-stranded cDNA (dscDNA), and incorporates dUTP into the second cDNA strand. cDNA generation is followed by end repair to generate blunt ends, A-tailing, adaptor ligation and PCR amplification. Different adaptors were used for multiplexing samples in one lane. Sequencing was performed on Illumina NovaSeq 6000 for PE 2×50 run. Data quality check was done on Illumina SAV. Demultiplexing was performed with Illumina Bcl2fastq v2.19.1.403 software.

### RNA Sequencing Analysis

Fastq files were run through FastQC and MultiQC to check the quality of raw sequence data (42, 43). A custom reference genome was generated by combining the Genome Reference Consortium Human build 38 (GRCh38/hg38) and the HIV-1 genome specific to J-Lat 10.6 (7) before aligning the reads to the genome through STAR (44). Aligned reads were then normalized through the DESeq2 package in R (45). R package EnhancedVolcano was used to visualize specific gene subsets (46). Gene lists were sorted and filtered for log2FoldChange above 2. Upregulated and downregulated gene lists were submitted to Metascape [https://metascape.org] for drug specific and cross comparison analyses (47). Top 500 up/downregulated genes were taken for drug specific Metascape analysis. Top 1000 up/downregulated genes were taken for drug cross comparison Metascape analysis. Gene set enrichment analysis (GSEA) was performed to identify the statistically significant hallmark gene sets from each drug treatment compared to untreated cells (48, 49).

### Immunofluorescence Microscopy

U2OS cells were plated on glass coverslips in 6-well tissue culture-treated plates (Greiner Bio-one) overnight and then switched into plain or drugged media consisting of either 5.0 ng/mL PMA, 50.0 µM Etoposide, 5.0 µM NSC95397, or 10.0 µL DMSO for 24 hours. Cells were then fixed in 4% PFA for 15 minutes and permeabilized with 1% Triton-X 100 in PBS for 5-10 min at 4°C. Cells were then blocked with immunofluorescence (IF) block buffer (3% BSA in PBS) for 1 hour and incubated with primary antibodies: rabbit anti-γH2A.x (Cell Signaling 9718, 1:2000), rabbit anti-RPA32 (GeneTex GTX113004 1:2000), or rabbit anti-53BP1 (Cell Signaling S1778, 1:2000) overnight at 4°C. Coverslips were washed three times with PBS then incubated with secondary antibodies (Alexa Fluor 488 or 594) and Hoechst 33342 (1:2000) for 1 hour. Coverslips were then washed three times with PBS and mounted onto slides with ProLong™ Gold Antifade Mountant (Invitrogen) and sealed when dried. Images were obtained with a ZEISS LSM 980.

### Data Availability

Data description: RNA sequencing data of J-Lat 10.6 cells untreated, or treated with PMA, SAHA, or NSC95397 Repository: Gene Expression Omnibus – NCBI Access Link: https://www.ncbi.nlm.nih.gov/geo/query/acc.cgi?acc=GSE241617

**Supplemental Figure 1.**
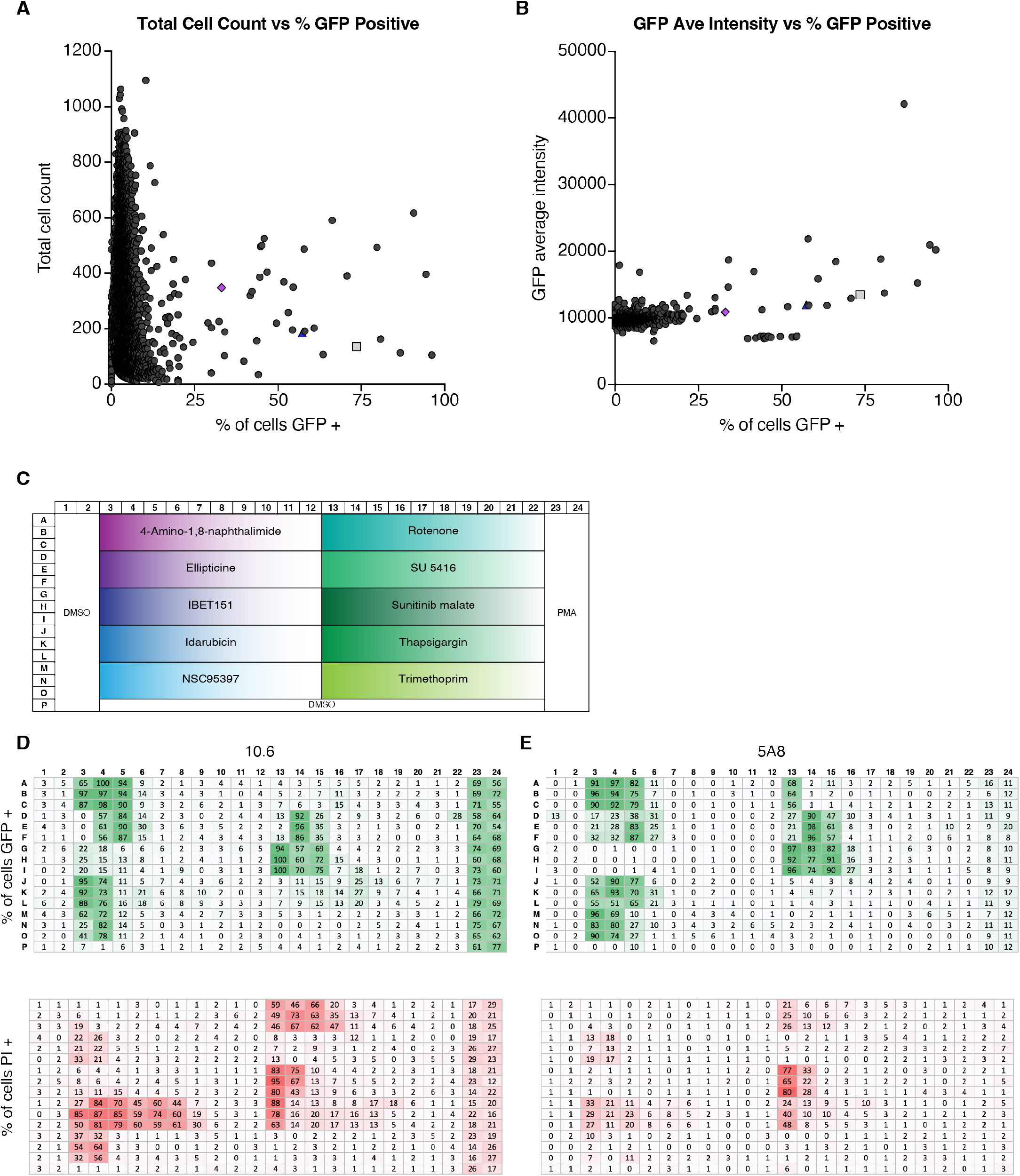
NSC95397 and IBET151 were identified as HIV-1 LRAs of interest from a medium-throughput screen. **A-B)** The raw screen data of total cell count or GFP intensity vs percent of cells GFP positive. The grey square and purple diamond are two known latency reversal agents, PMA and IBET151, respectively, and the blue triangle is a previously unknown LRA, NSC95397. **C)** Example plate map of the EC_50_ screen. Each compound was screened in triplicate and across the plate is a 4-fold concentration decrease. The first two columns and last row were treated with the vehicle control, DMSO, and the last two columns were treated with the positive control, PMA. **D-E)** The percent of GFP positive cells, top green plate, and dead PI positive cell, lower red plate, measured during our EC_50_ titration experiment in J-Lat 10.6 **(D)** and 5A8 cells **(E)**. Related to Figure 1.

**Supplemental Figure 2.**
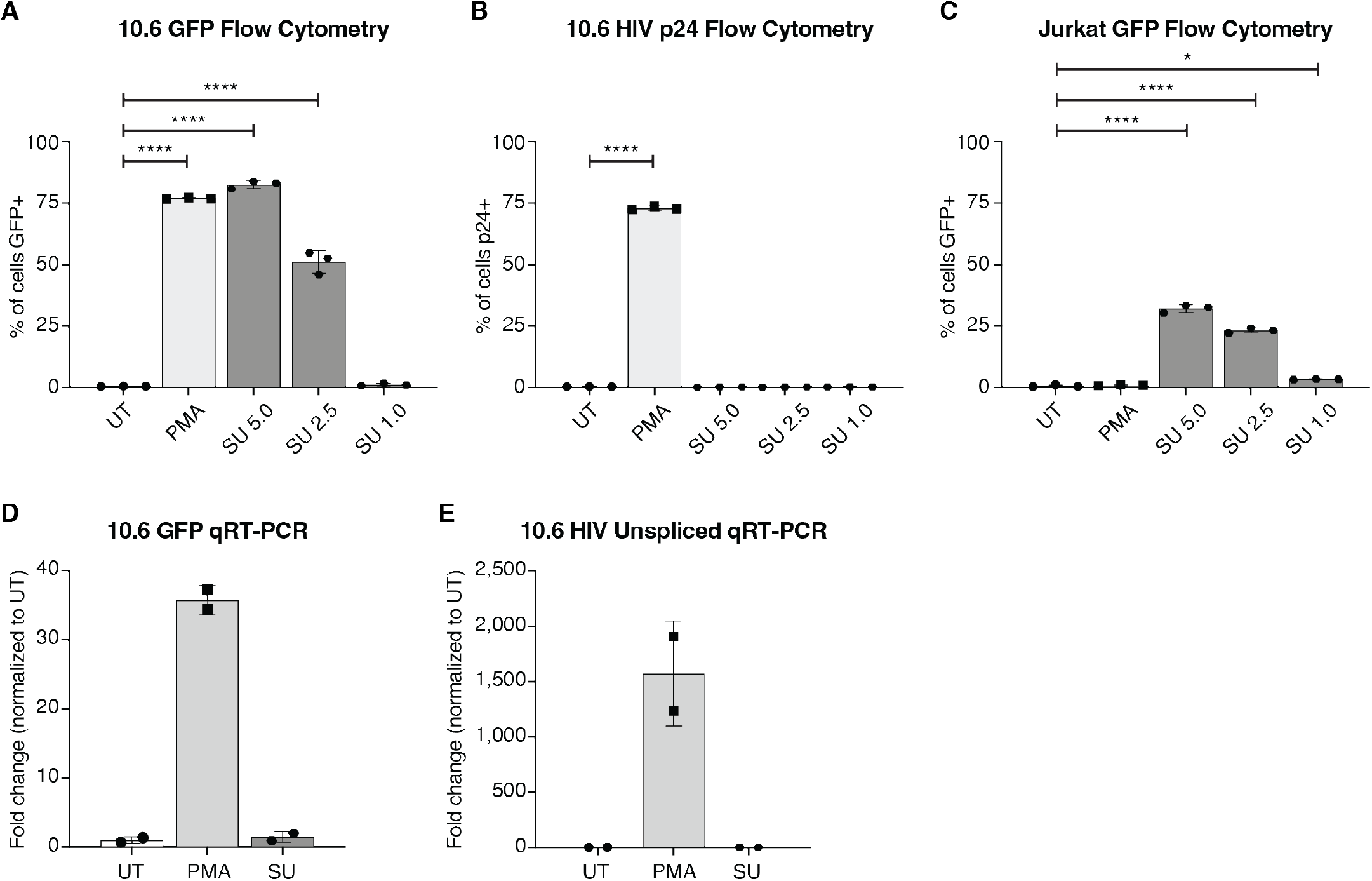
Sunitinib is one of a subset of false positives identified in a J-Lat screen for LRAs. J-Lat 10.6 or parental Jurkat cells were untreated or treated with 5.0 ng/mL PMA (positive control), or a titration of sunitinib (SU) from 5.0 to 1.0 μM. **A-B)** Bar graphs showing GFP (**A**) or intracellular HIV-1 p24 (**B**) measured via flow cytometry in J-Lat 10.6 cells. **C)** A bar graph of GFP measured via flow cytometry in treated Jurkat cells. **D-E)** Bar graphs showing GFP (**D**) or unspliced HIV-1 RNA transcripts (**E**) measured via qRT-PCR. Related to Figure 2

**Supplemental Figure 3.**
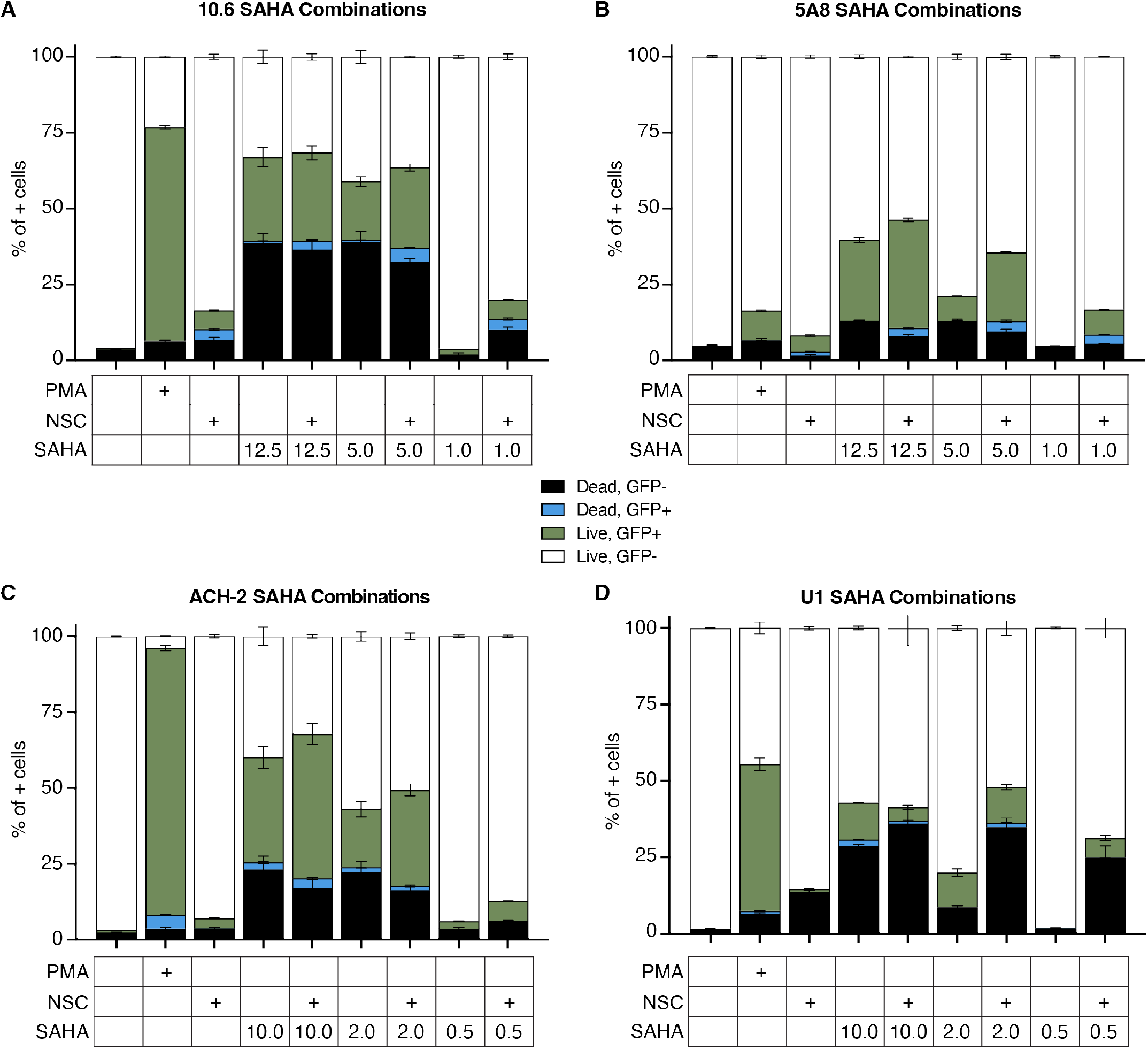
NSC95397 increases HIV-1 reactivation and cytotoxicity in combination with SAHA. Stacked graphs show the percentage of cells from the full population that are either dead (ghost dye positive) and GFP- (black), dead and GFP+ (blue), live and GFP+ (green), or live and GFP- (white) stacked to a total of 100%. **A-B)** J-Lat 10.6 (**A**) or 5A8 (**B**) cells were untreated or treated with 5.0 ng/mL PMA, 1.5 μM NSC95397 (NSC), a titration of SAHA from 12.5 to 1.0 μM, or a titration of SAHA with NSC held constant at 1.5 μM. **C-D)** ACH-2 (**C**) or U1 (**D**) cells were untreated or treated with 5.0 ng/mL (**C**) or 20.0 ng/mL PMA (**D**), 2.5 μM NSC, a titration of SAHA from 10.0 to 0.5 μM, or a titration of SAHA with NSC held constant at 2.5 μM. Related to Figure 3.

**Supplemental Figure 4.**
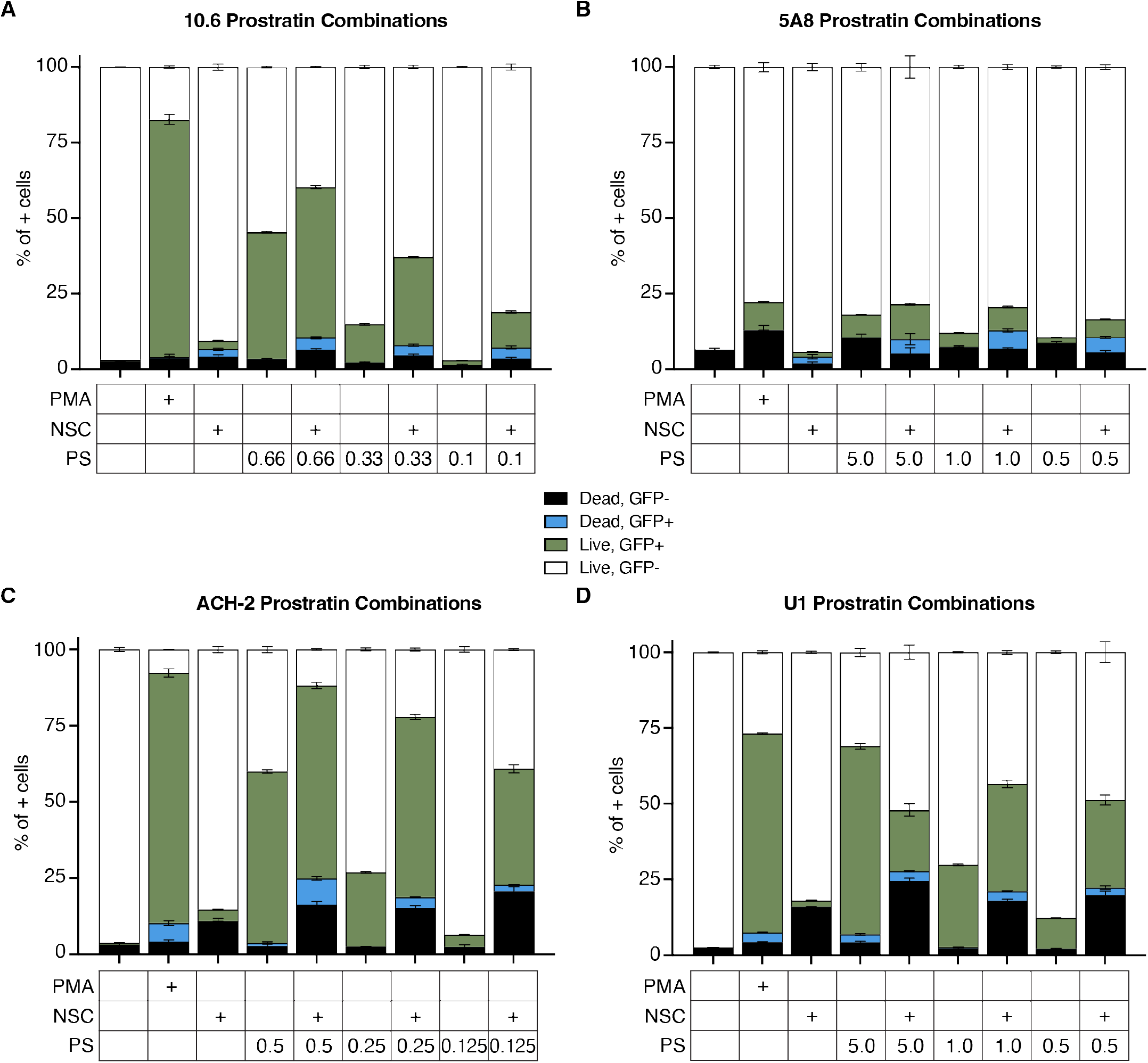
NSC95397 increases HIV-1 reactivation and cytotoxicity in combination with prostratin. Stacked graphs show the percentage of cells from the full population that are either dead (ghost dye positive) and GFP- (black), dead and GFP+ (blue), live and GFP+ (green), or live and GFP- (white) stacked to a total of 100%. **A-B)** J-Lat 10.6 (**A**) or 5A8 (**B**) cells were untreated or treated with 5.0 ng/mL PMA, NSC95397 (NSC) at 1.5μM, a titration of prostratin (PS) from 0.66 to 0.1 μM (**A**) or 5.0 to 0.5 μM (**B**), or a titration of PS with NSC held constant at 1.5 μM. **C-D)** ACH-2 (**C**) or U1 (**D**) cells were untreated or treated with 5.0 ng/mL (**C**) or 20.0 ng/mL PMA (**D**), 2.5 μM NSC, a titration of PS from 0.5 to 0.125 μM (**C**) or 5.0 to 0.5 μM (**D**), or a titration of PS with NSC held constant at 2.5 μM. Related to Figure 4.

**Supplemental Figure 5.**
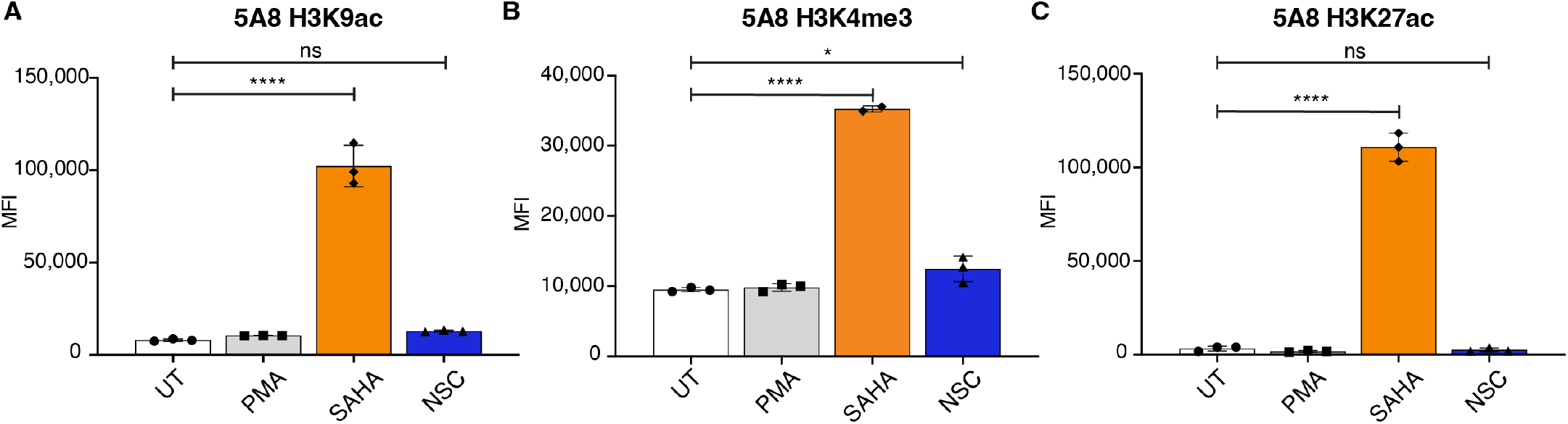
NSC95397 does not change global histone modifications for open chromatin. **A-C)** J-Lat 5A8 cells were untreated or treated with 5.0 ng/mL PMA, 25.0 μM SAHA (positive control), or NSC95397 (NSC) at 2.5 μM. Bar graphs show H3K9ac (**A**), H3K4me3 (**B**), or H3K27ac (**C**) MFI measured by flow cytometry. Bar graphs show GFP MFI measured via flow cytometry. Results were analyzed with a one-way ANOVA with Turkey’s multiple comparisons tests; ns = not significant, * = p ≤ 0.0332, ** = p ≤ 0.0021, *** = p ≤ 0.0002, **** = p ≤ 0.0001. Related to Figure 5.

**Supplemental Figure 6.**
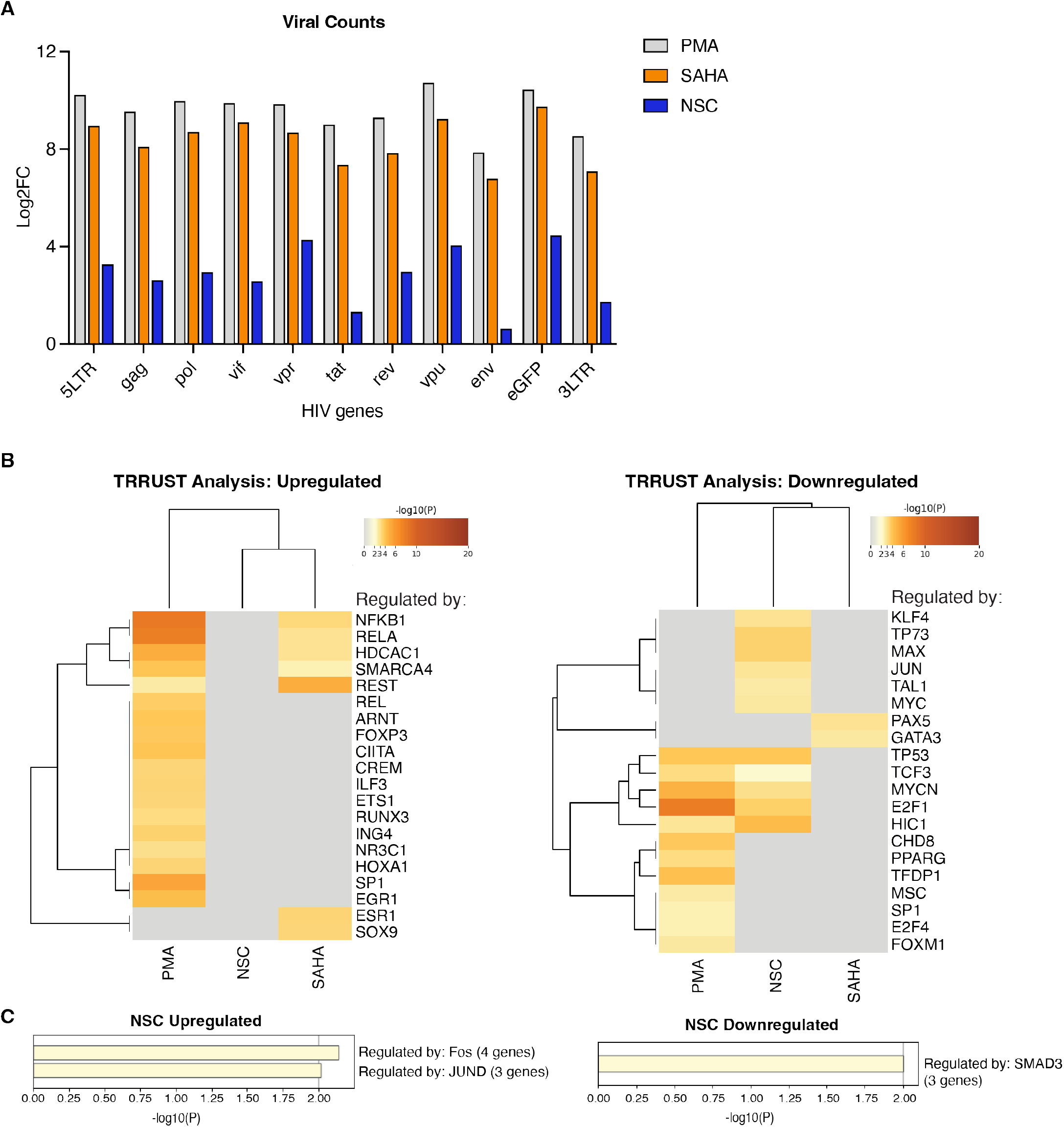
NSC95397 has a minimal effect on global transcription. J-Lat 10.6 cells were untreated or treated with 5.0 ng/mL PMA, 25.0 μM SAHA, or 5.0 μM NSC95397 (NSC) and used for bulk RNA sequencing in four biological replicates. **A)** Bar graph showing log2 fold change of HIV-1 associated reads normalizing cells treated with PMA (grey), SAHA (orange), or NSC (blue) against untreated cells. **B-C)** Gene ontology using Metascape for TRRUST analysis of upregulated (left) and downregulated (right) genes for all three treatments (**B**) and NSC alone (**C**). Related to Figure 6 and Supplemental Figure 7.

**Supplemental Figure 7.**
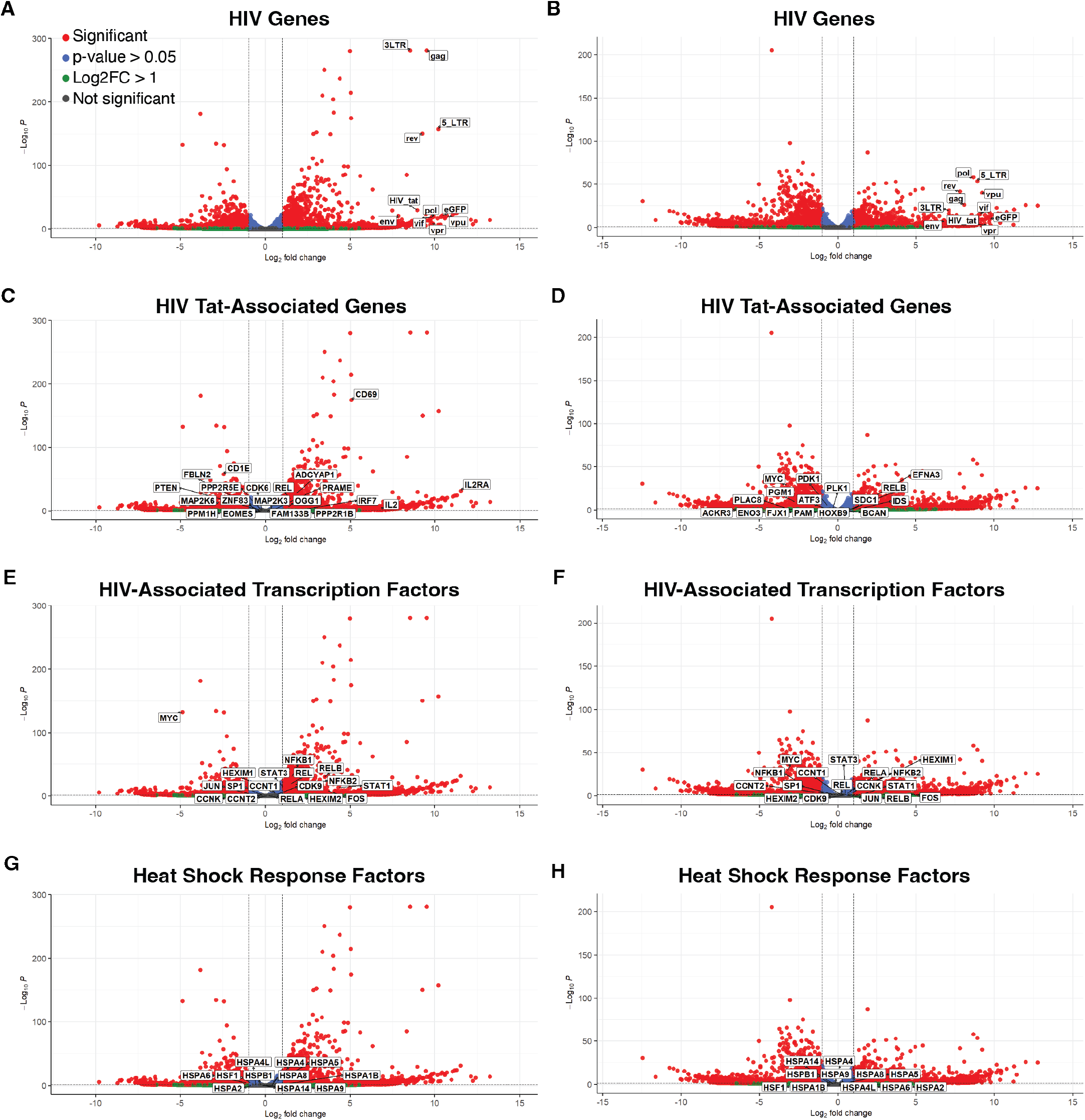
PMA and SAHA change latency modulating pathways. J-Lat 10.6 cells were untreated or treated with 5.0 ng/mL PMA or 25.0 μM SAHA and used for bulk RNA sequencing in four biological replicates. **A-B)** Volcano plot highlighting HIV-associated reads in cells treated with PMA (**A**) or SAHA (**B**). **C-D)** Volcano plots highlighting HIV-associated transcription factors in cells treated with PMA (**C**) or SAHA (**D**). **E-F)** Volcano plots highlighting selected HIV Tat-modulated genes in cells treated with PMA (**E**) or SAHA (**F**). **G-H)** Volcano plots highlighting key heat shock response genes in cells treated with PMA (**G**) or SAHA (**H**). Cut-offs are drawn for log2 fold change above 1 and p-value greater than 0.05 where reads are separated to non-significant and non-enriched (gray), non-significant with log2 fold change above 1.0 (green), significant and log2 fold change below 1.0 (blue), or significant and log2 fold change above 1.0 (red). Related to Figure 6 and Supplemental Figure 6.

**Supplemental Figure 8.**
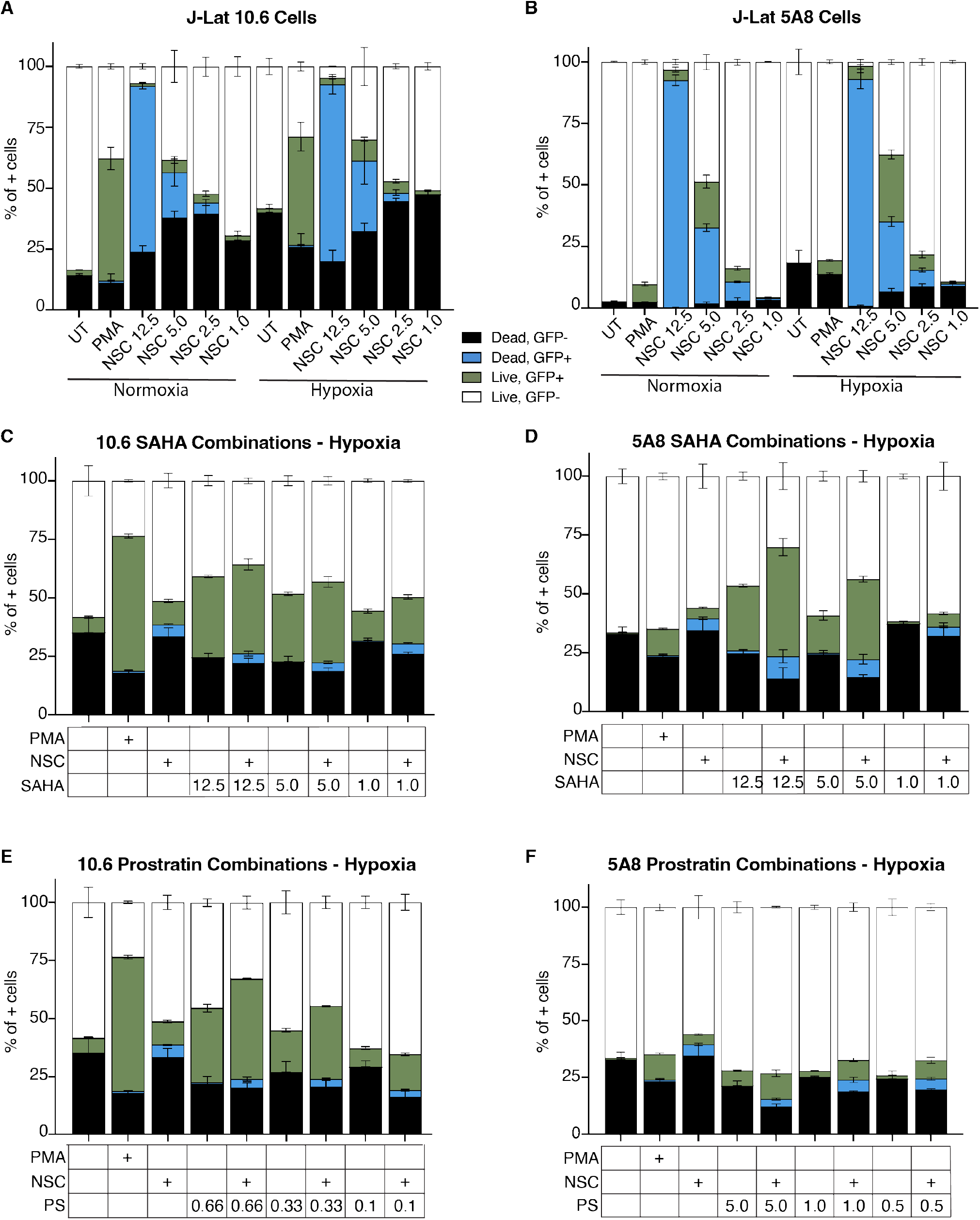
NSC95397 alone and in combination with LRAs increases HIV reactivation and cytotoxicity in cells under hypoxia. **A-F)** Stacked graphs show the percentage of cells from the full population that are either dead (ghost dye positive) and GFP- (black), dead and GFP+ (blue), live and GFP+ (green), or live and GFP- (white) stacked to a total of 100%. Bar graphs show the percent of cells that are GFP+ from the full population. **A-B)** J-Lat 10.6 (**A**) or 5A8 (**B**) cells were untreated or treated with 5.0 ng/mL PMA (positive control), or a titration of NSC95397 (NSC) from 5 μM to 1.0 μM **C-D)** J-Lat 10.6 (**C**) or 5A8 (**D**) cells were untreated or treated with 5.0 ng/mL PMA (positive control), NSC95397 (NSC) at 1.5 μM, a titration of SAHA from 12.5 to 1.0 μM, or a titration of SAHA with NSC held constant at 1.5 μM. **E-F)** J-Lat 10.6 (**E**) or 5A8 (**F**) cells were untreated or treated with 5.0 ng/mL PMA, NSC at 1.5 μM, a titration of prostratin (PS) from 0.66 to 0.1 μM (**E**) or 5.0 to 0.5 μM (**F**), or a titration of PS with NSC held constant at 1.5 μM. Results were analyzed with a one-way ANOVA with Turkey’s multiple comparisons tests; ns = not significant, * = p ≤ 0.0332, ** = p ≤ 0.0021, *** = p ≤ 0.0002, **** = p ≤ 0.0001. Related to Figure 6 and Supplemental 7.

